# Dysregulation of Hippo Signaling Pathway as a Convergent Mechanism Underlying Choroid Plexus Defects in Bipolar Disorder

**DOI:** 10.64898/2026.04.10.717745

**Authors:** Jonghun Kim, Mu Seog Choe, Kutlu Kaya, Yue Hu, Kenneth K. Ng, Caihong Qiu, Mei Zhong, Woo Sub Yang, Ferdi Ridvan Kiral, Cynthia Lo, Michael Scandura, Hyuk-Jin Cha, Jeffrey R. Gruen, Monkol Lek, Elizabeth A. Jonas, Hilary P. Blumberg, In-Hyun Park

## Abstract

Bipolar disorder (BD) is a prevalent and highly heritable psychiatric condition. Developmental mechanisms are implicated but the specific molecular origins remain unclear. The choroid plexus (ChP), which regulates cerebrospinal fluid (CSF) and brain homeostasis, has been implicated in BD, but its role is poorly understood. Here, we identify aberrant expansion of ChP in human cortical (hCOs) and ChP organoids (hChPOs) derived from, and *in vivo* structural magnetic resonance imaging (sMRI) scans of, individuals with BD compared to healthy comparison individuals. Single-cell transcriptomics revealed a lineage bias toward ChP fate in BD hCOs, accompanied by structural enlargement observed in both BD hChPOs and ChP in sMRI scans from individuals with BD. Comprehensive transcriptomic profiling implicates early hyperactivation of the Hippo signaling pathway in neural progenitor cells as a convergent mechanism driving ChP overgrowth. Whole-genome and genome-wide association study analyses highlight recurrent genetic variants in Hippo regulators, including *STK4* and *YAP1*, suggesting genetic priming. BD hChPOs exhibit disrupted epithelial junctions and altered *in vitro* CSF secretion. These findings position the ChP as a genetically and developmentally predisposed to pathology in BD and highlight organoids as a platform for translational discovery in diagnostics and therapeutics.

## Introduction

Bipolar disorder (BD) affects about 4% of the population^1^. Its characteristic manic and depressive episodes are associated with immense suffering, disability, elevated medical morbidity, and premature mortality due to both medical complications and high suicide rates^2,3^. Structural brain abnormalities have been observed across multiple regions^4,5^. Increased lateral ventricular size was one of the first and remains one of the most consistent findings^6^. In addition, genome-wide association studies (GWAS) have identified multiple genetic risk loci associated with BD^7–9^, highlighting a strong genetic contribution to disease susceptibility. Despite its clinical severity and well-established heritability, the molecular and cellular mechanisms underlying brain differences in BD remain poorly understood.

The choroid plexus (ChP) is a specialized brain region responsible for cerebrospinal fluid (CSF) production and homeostasis, affecting ventricular size, and has recently been implicated in BD. Neuroimaging studies have shown ChP enlargement in individuals with BD^10–13^, yet the mechanisms underlying this structural expansion remain largely unknown. Traditional *in vivo* and postmortem approaches have provided limited insight into the cellular and developmental origins of pathological changes in brain development. To address these challenges, brain organoids generated from human induced pluripotent stem cells (iPSCs) provide a robust platform for recapitulating the genetic and cellular features of brain disorders in a human-specific context^14,15^.

Here, we demonstrate ChP enlargement in *in vitro* ChP model, ChP organoids (hChPOs), as well as in *in vivo* neuroimaging data from the individuals with BD. Transcriptomic profiling of BD organoids reveals dysregulation of the Hippo signaling pathway as a convergent mechanism. These findings are further corroborated by the enrichment of Hippo-related genes in BD GWAS loci and the rare variants observed in our whole-genome sequencing (WGS) data from individuals with BD. Moreover, BD hChPOs exhibit altered cell junction architecture and secrete *in vitro* CSF (iCSF) with distinct proteomic profile compared to control hChPOs **(Fig. 1a)**. Altogether, our study identifies ChP abnormalities as cellular and molecular features of BD and implicates Hippo signaling as a convergent pathway underlying these changes. By integrating neuroimaging, ultrastructural, genomic, transcriptomic, and proteomic analyses, our work provides a unified framework for understanding ChP dysfunction in BD.

**Fig. 1:**
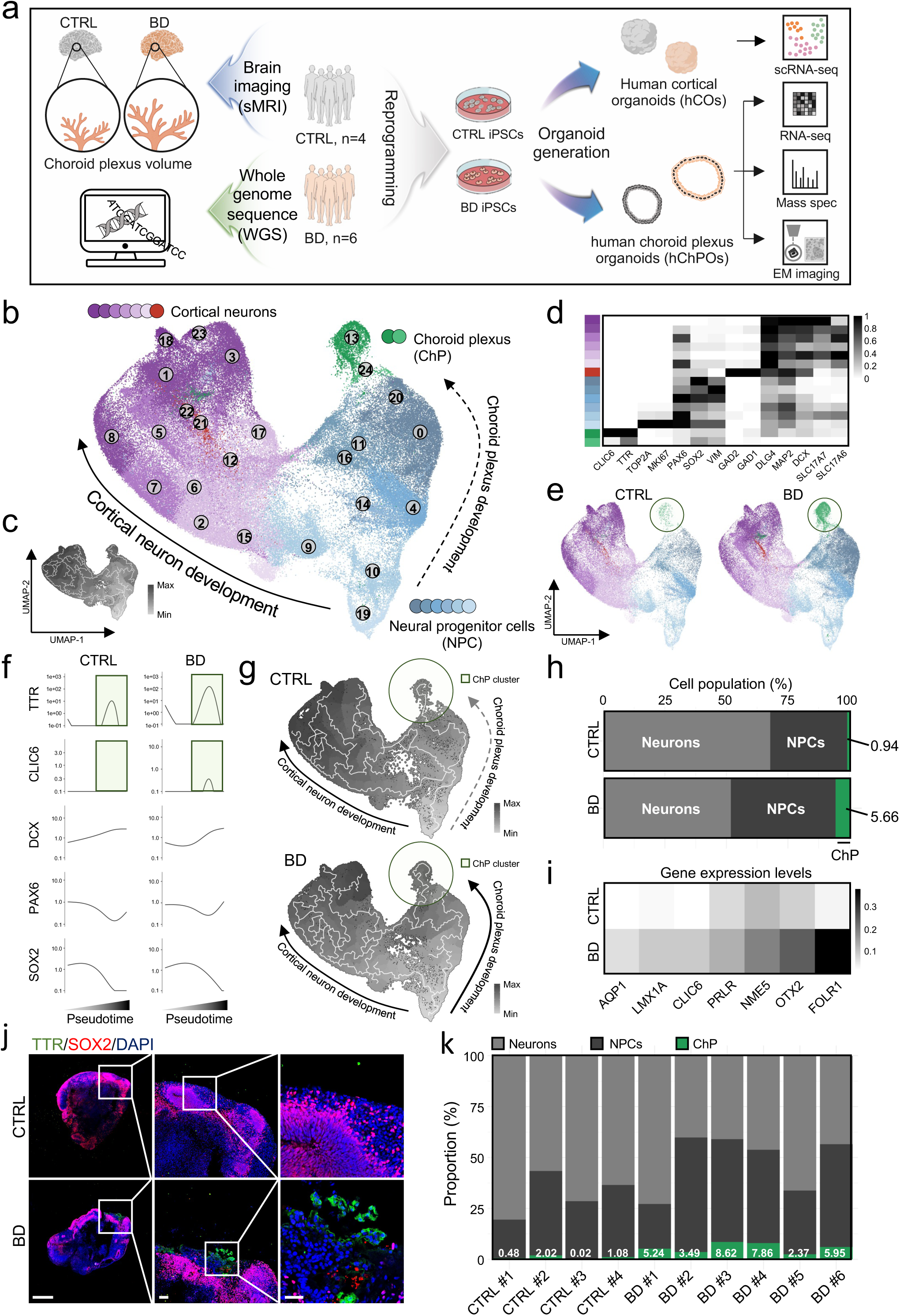
Single-Cell transcriptomics reveals ChP fate bias in BD hCOs. **a**, Schematic overview of the experimental workflow used in this study. **b**, UMAP projection of all cells from Day 75 hCOs derived from four control and six BD iPSCs. **c**, Reconstruction of developmental trajectories through pseudotime analysis. **d**, Heatmap illustrating cell type-specific expression profiles of selected marker genes. **e**, UMAP plots comparing cellular landscapes between control and BD hCOs. **f**, Temporal expression dynamics of ChP markers (*TTR*, *CLIC6*) and neuronal lineage markers (*DCX*, *PAX6*, *SOX2*) along pseudotime. **g**, Pseudotime-inferred developmental trajectories in control and BD hCOs. **h**, Comparison of cellular compositions between control and BD hCOs. **i**, Expression levels of ChP-specific markers. **j**, Immunofluorescence images of whole hCOs at Day 75 stained for TTR and SOX2. Nuclei are counterstained with DAPI. The white box indicates the magnified region shown on the right. Scale bars, 500 µm (left), 100 µm (middle and right). **k**, Cell type compositions across individual hCOs. Ctrl, control; NPC, neural progenitor cells; ChP, choroid plexus.

### Lineage Bias Toward a ChP Fate in BD

Recent genetic and neuroimaging studies suggest that BD may originate from subtle perturbations during early neurodevelopment^16,17^. To investigate the overall differences in neurodevelopmental trajectories and cortical cell-type composition in BD, we first generated human iPSCs from adult females with BD (*n* = 6) and healthy control adult females (*n* = 4). Peripheral blood mononuclear cells (PBMCs) were isolated from these donors and reprogrammed to generate iPSC lines. All 10 iPSC lines expressed core pluripotency markers, including *OCT4*, *NANOG*, *SOX2*, and *REX1*, as validated by RT-qPCR **(Extended Data Fig. 1a)**. Consistent with these results, immunohistochemistry for OCT4 and NANOG showed uniform nuclear staining across all iPSC lines, further confirming their pluripotent identity **(Extended Data Fig. 1b)**.

Next, we generated hCOs from the 10 iPSC lines. All 10 hCOs exhibited typical neuroepithelial features, with no discernible differences observed under bright-field microscopy **(Extended Data Fig. 2)**. We then performed single-cell RNA sequencing (scRNA-seq) at differentiation day 75, a stage at which the major cortical lineages are established and both neural progenitor cells (NPCs) and mature neurons coexist. scRNA-seq profiles from all hCOs were integrated, resolving 25 transcriptionally distinct clusters **(Extended Data Fig. 3a).** Marker gene expression clearly segregated these clusters into three principal lineages, NPCs, neurons, and ChP, based on canonical neuronal development markers (*PAX6*, *SOX2*, *DCX*, *SLC17A7*) and ChP-specific markers (*TTR*, *CLIC6*) **(Extended Data Fig. 3b,c)**. UMAP embedding revealed a bifurcated developmental landscape comprising a cortical neuron trajectory and a ChP trajectory **(Fig. 1b,c),** with lineage-defining markers faithfully demarcating cluster borders **(Fig. 1d).**

When the dataset was split by clinical group, the ChP clusters were significantly larger in BD hCOs **(Fig. 1e)**. To further investigate dynamic lineage decisions, we performed pseudotime analysis. While the neuronal development markers (*DCX*, *PAX6*, *SOX2*) displayed similar developmental trajectories, ChP markers *TTR* and *CLIC6* showed sharp increases in BD hCOs **(Fig. 1f).** Group-specific trajectory reconstruction confirmed an expanded ChP population and a more intricate developmental path in BD hCOs **(Fig. 1g).** Quantitatively, ChP population constituted only 0.94% of the control atlas but 5.66% of the BD atlas, representing a more than 5.5-fold increase **(Fig. 1h)**, accompanied by broad upregulation of additional ChP markers **(Fig. 1i)**. Finally, we asked whether the transcriptomic expansion of the ChP lineage manifested at the tissue level and independently validated the enlarged ChP structure in BD hCOs using TTR immunohistochemistry **(Fig. 1j)**.

To exclude the possibility that the observed ChP expansion resulted from integration or batch-related artifacts, we reanalyzed each hCO separately **(Extended Data Fig. 4a)**. The increase in ChP population was consistently observed across individual BD hCOs **(Fig. 1k)**. At the single-sample level, TTR expression was consistently elevated in all BD hCOs compared to control hCOs **(Extended Data Fig. 4b)**, with other ChP markers, including *FOLR1*, *OTX2*, and *NME5*, showing a similar upward trend **(Extended Data Fig. 4c)**. When UMAP embeddings were split by sample, the ChP cluster appeared visibly enlarged in all BD hCOs **(Extended Data Fig. 4d)**. Moreover, pseudotime trajectories confirmed that *CLIC6* and *TTR* expression remained elevated across all BD hCOs relative to control hCOs **(Extended Data Fig. 4e)**. Together, these individual-sample analyses demonstrate that the pronounced ChP fate bias in BD hCOs represents a robust biological phenomenon rather than a technical artifact.

Collectively, these data suggest that BD hCOs exhibit a consistent bias toward the ChP lineage during early cortical development, implicating aberrant ChP expansion as a previously unrecognized contributor to BD pathogenesis.

### ChP Enlargement in Individuals with BD

To evaluate the clinical relevance of scRNA-seq findings, we analyzed structural magnetic resonance imaging (sMRI) scan data from the individuals whose iPSCs were derived and used to generate hCOs. Structural scans were available for all six individuals with BD and three out of the four control individuals. We assessed total, left, and right ChP and lateral ventricle (LV) volumes. ChP total volume was higher in individuals with BD, with a moderate effect size Cohen’s *d* = 0.4. Although the comparison to the control sample was not significant, left ChP in individuals with BD were larger than in controls with Cohen’s *d* = 0.49. **(Extended Data Fig. 5a-c)**. LV expansion has long been reported in BD^6^. We observed total LV volume enlargement in the individuals with BD (Cohen’s *d* = 0.36 for total LV volume), with similar effects for the left and right LV, although the comparison to the controls was not significant **(Extended Data Fig. 5d-f)**. Taken together, these data suggest that ChP alterations identified in hCOs may extend to ChP and LV alterations in the brains of adults with BD.

### BD hChPOs Exhibit Increased Size

We next generated human choroid plexus organoids (hChPOs) from the control and BD iPSCs using a previously established protocol^18^ with slight modifications **(Fig. 2a)**. Cyst-like structures emerged by day 15 of differentiation, indicating early ChP-like tissue organization. By day 35, the cuboidal epithelial cells lining the cyst-like structures appeared well-defined and substantially enlarged **(Fig. 2b)**. By day 80, cortical regions had diminished, and most organoids had developed into fluid-filled cysts **(Fig. 2c).**

**Fig. 2:**
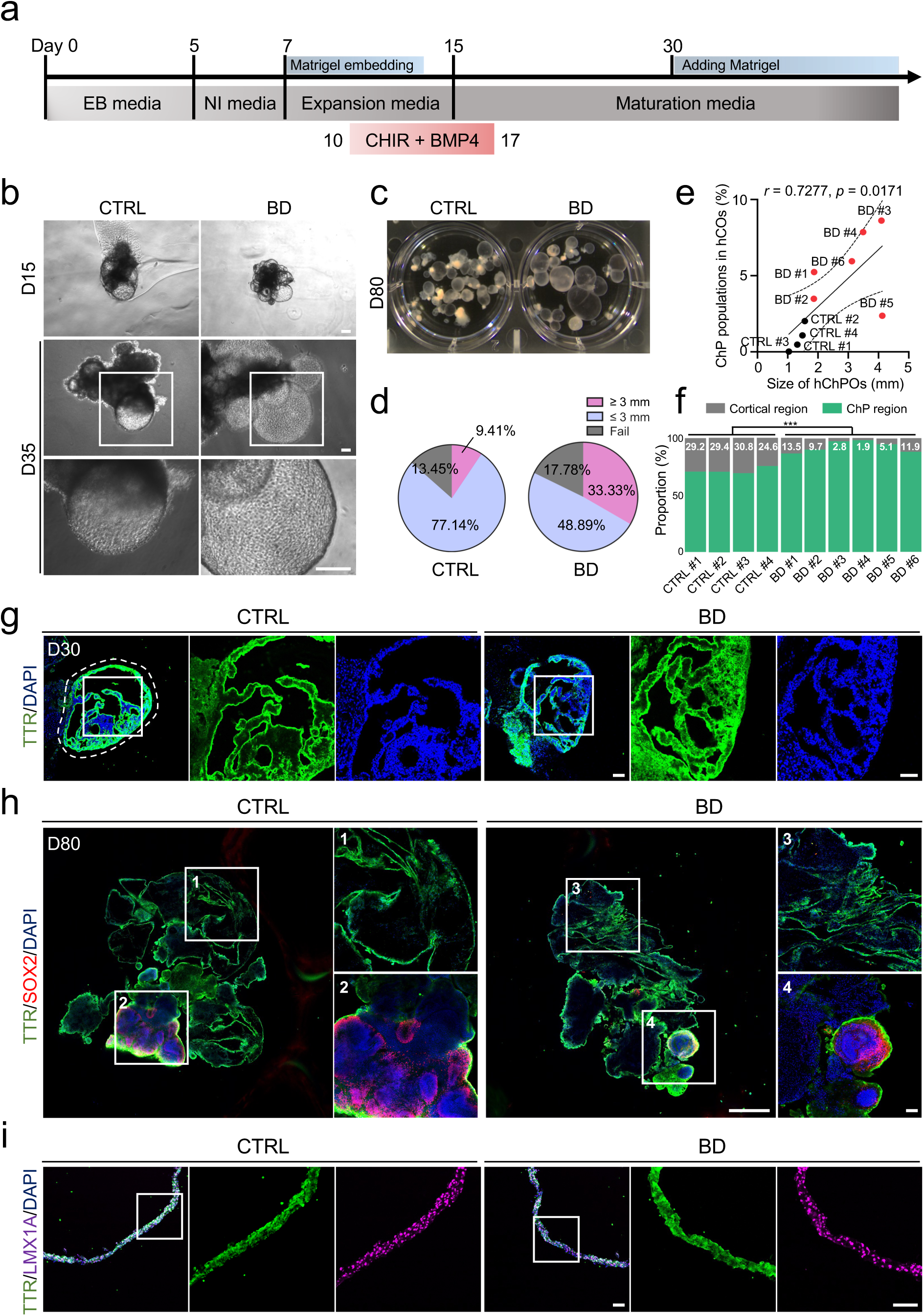
BD hChPOs are enlarged. **a**, Schematic of the protocol for generating hChPOs from human iPSCs. **b**, Representative images of hChPOs at Day 15 (top), Day 35 (middle), and a high-magnification view showing the cuboidal organization of the ChP epithelium (bottom). Scale bars, 100 µm. **c**, Representative images of hChPOs exhibiting fluid-filled cystic morphology at Day 80. **d**, Pie chart showing the proportional distribution of sizes in control (*n* = 59) and BD (*n* = 61) hChPOs. **e**, Correlation between ChP population abundance in hCOs and average hChPO size across samples (*r* = 0.7277, *p* = 0.0171). **f**, Regional composition of ChP and cortical domains across all hChPOs (*n* = 5-10 per sample). **g**, Immunofluorescence images of TTR expression in hChPOs at Day 30. Nuclei are counterstained with DAPI. The white box indicates the magnified region shown on the right. Scale bars, 100 µm. **h**, Whole-mount immunofluorescence images of control and BD hChPOs at Day 80 stained for TTR and SOX2. Nuclei are counterstained with DAPI. White boxes indicate the magnified regions shown in panels 1–4, with panels 1 and 3 highlighting ChP regions and panels 2 and 4 highlighting cortical regions. Scale bars, 500 µm (main), 100 µm (1–4). **i**, Immunofluorescence images of TTR and LMX1A expression in hChPOs at Day 80. Nuclei are counterstained with DAPI. The white box indicates the magnified region shown on the right. Scale bars, 100 µm. Statistical analysis was performed using Pearson’s correlation test (e) or a two-tailed Welch’s t-test (f). \*\*\**P* < 0.001.

We then compared organoid size and structural composition between control and BD hChPOs. Quantitative analyses were restricted to ChP regions in hChPOs to assess ChP-specific alterations. The BD group exhibited a higher proportion of large-sized organoids (≥ 3 mm) relative to controls **(Fig. 2d and Extended Data Fig. 6a)**. A strong positive correlation (r = 0.73) was identified between the proportion of ChP populations in hCOs, as determined by scRNA seq, and the size of hChPOs across both control and BD groups **(Fig. 2e and Supplementary Table 1)**, suggesting a robust relationship between ChP specification and growth across two independent organoid models. We further analyzed the relative proportions of cortical and ChP regions within each organoid to compare structural composition between control and BD hChPOs. In line with our previous transcriptomic findings, BD hChPOs exhibited a marked expansion of the ChP region relative to control hChPOs **(Fig. 2f)**, driven by increases in both size and proportion. These findings suggest that BD hChPOs undergo an expansion of ChP lineages, resulting in both an increase in overall organoid size and a relative dominance of the ChP region. To characterize control and BD hChPOs, we performed immunohistochemical analysis. At day 30, TTR-positive cells were arranged in cyst-like formations, suggestive of early ChP morphogenesis **(Fig. 2g)**. As the hChPOs matured, the number of TTR-positive cells progressively increased. By day 80, TTR was expressed in the majority of cells, with this trend more pronounced in BD hChPOs **(Fig. 2h)**. Additionally, expression of the ChP marker LMX1A was observed in both control and BD hChPOs, indicating proper ChP differentiation in both groups **(Fig. 2i and Extended Data Fig. 6b)**.

Altogether, these data suggest that although control and BD hChPOs share key molecular and structural features, BD hChPOs exhibit an aberrant expansion in size, potentially reflecting disease-associated alterations in ChP development.

### Hyperactivation of the Hippo Signaling Pathway Drives ChP Overgrowth in BD

Having documented the marked expansion of the ChP lineage, we next sought to uncover the molecular mechanisms underlying this overgrowth by systematically profiling hChPOs. To investigate the molecular basis of ChP overgrowth, we performed RNA-seq on the control and BD hChPOs **(Fig. 3a)**. Principal-component analysis (PCA) positioned the two clinical groups in opposite quadrants **(Fig. 3b)**, and a hierarchical clustering heatmap reproduced this sharp separation **(Extended Data Fig. 7a)**, underscoring a fundamental transcriptomic divergence. Before examining signaling pathways, we first asked whether the overgrowth of BD hChPOs could be explained by altered CSF production. To address this, we examined the expression of key ion channel and transporter genes involved in CSF secretion, including *AQP1*, *CA2*, and *SLC4A5*. Expression levels of these genes did not differ between control and BD hChPOs **(Extended Data Fig. 7b)**, suggesting that changes in CSF production are unlikely to be the primary driver of ChP expansion.

**Fig. 3:**
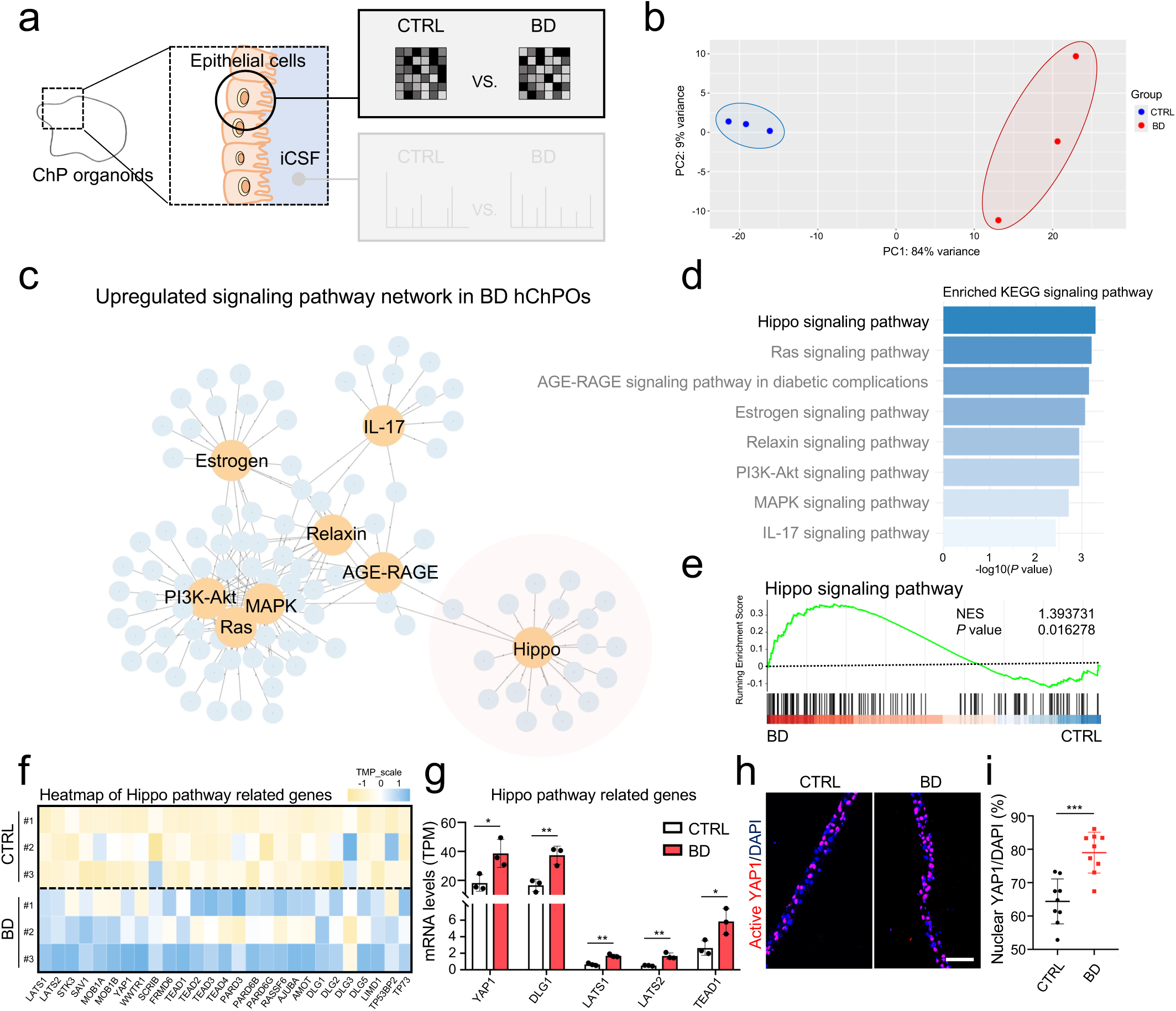
Transcriptomic profiling identifies aberrant Hippo pathway in BD hChPOs. **a**, Schematic outlining the transcriptomic analysis of control and BD hChPOs at Day 80. **b**, PCA illustrating transcriptomic differences between control and BD hChPOs (n = 3 per group, biological replicates). **c**, Network of enriched KEGG pathways, highlighting signaling pathways upregulated in BD hChPOs. The Hippo signaling pathway is highlighted with a red circle. **d**, Enrichment plot of upregulated signaling pathways in BD hChPOs, ranked by statistical significance (–log_10_*P*). **e**, GSEA performed on the Hippo signaling pathway in control and BD hChPOs. **f**, Heatmap showing expression levels of Hippo signaling pathway genes in control and BD hChPOs. **g**, Expression levels of core components of the Hippo signaling pathway, based on TPM values (*n* = 3 per group). **h**, Immunofluorescence images of active YAP1 expression in control and BD hChPOs. Scale bars, 100 µm. **i**, Quantification of active YAP1-positive cells (*n* = 9 per group). All data are presented as mean ± s.d. Statistical analysis was performed using a two-tailed Student’s t-test (g,i). \*\*\**P* < 0.001, \*\**P* < 0.01, \**P* < 0.05.

To comprehensively assess transcriptional alterations beyond transport function of ChP, we next carried out a global differential expression analysis between the two groups. Differential expression analysis followed by KEGG enrichment-based network highlighted the Hippo, MAPK, and PI3K-Akt signaling pathways **(Fig. 3c)**. Among dysregulated signaling pathways, Hippo signaling pathway emerged as the principal axis of transcriptional change **(Fig. 3d)**. Gene set enrichment analysis (GSEA) confirmed significant upregulation of the Hippo signaling pathway in BD hChPOs **(Fig. 3e)**. Core components, including *YAP1*, *TEAD1*, *LATS2*, and *WWTR1/TAZ*, were uniformly upregulated **(Fig. 3f)**, a trend further validated by quantitative analysis **(Fig. 3g)**. Consistent with pathway activation, immunohistochemistry for the active form of YAP1, a key transcriptional co-activator in the Hippo pathway, supported the transcriptomic data, showing substantially increased nuclear translocation in BD hChPOs **(Fig. 3h,i)**. In addition, canonical downstream targets of YAP/TAZ-mediated transcription, including *CYR61*, *AXL*, and *AMOTL2*, were markedly induced **(Extended Data Fig. 7c).**

Taken together, these data implicate aberrant activation of the Hippo signaling pathway, driven by elevated YAP1 activity, as a key driver of the pronounced ChP fate bias observed in BD organoids, thereby providing a mechanistic link between transcriptomic dysregulation and ChP expansion.

### Early Activation of the Hippo Signaling Pathway in BD Diverts NPCs toward a ChP Fate

To incorporate and further analyze temporal information based on the developmental trajectory, we revisited our hCO scRNA-seq data to map Hippo signaling pathway along single-cell developmental trajectories. This approach preserves lineage-specific timing and allows identification of when pathway divergence begins. In the ChP clusters, core components of the Hippo signaling pathway, such as *LATS2*, *YAP1*, and *TEAD1*, were markedly upregulated in BD hCOs **(Fig. 4a)**. This concordance supports the conclusion that the transcriptomic changes observed in bulk RNA-seq of BD hChPOs reflect ChP lineage-specific expression rather than artifacts arising from sample heterogeneity.

**Fig. 4:**
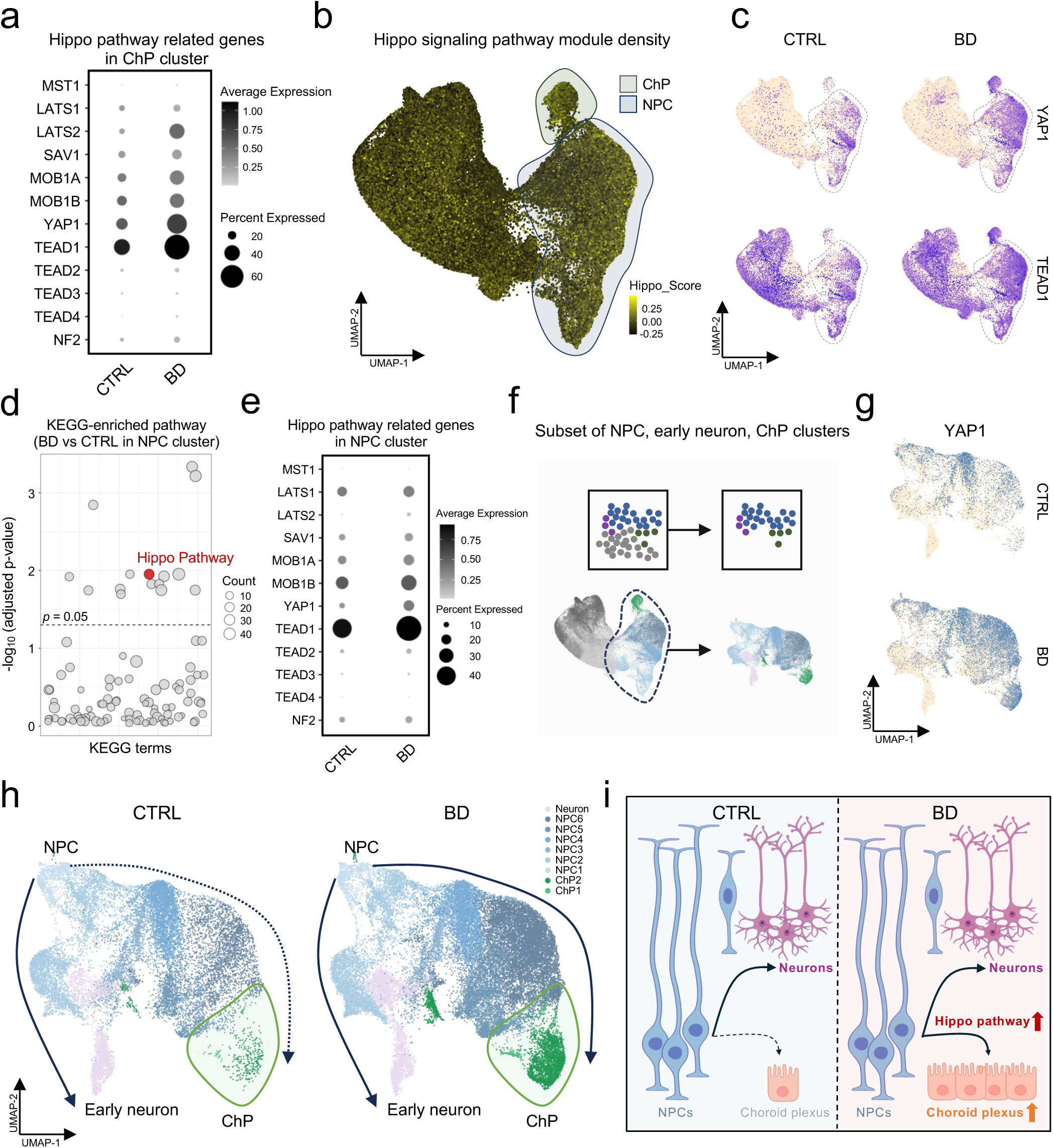
Early Hippo pathway activation in NPCs promotes ChP fate bias in BD hCOs. **a**, Dot plot showing expression levels of Hippo pathway-related genes in the ChP cluster. **b**, Module scoring of Hippo pathway activity across all clusters. **c**, Feature plots displaying expression of core Hippo signaling components, *YAP1* and *TEAD1*. **d**, KEGG enrichment analysis of the NPC cluster in control and BD hCOs, using a *P* value threshold of < 0.05. The Hippo signaling pathway is highlighted in red. **e**, Dot plot showing expression levels of Hippo pathway-related genes in the NPC cluster. **f**, Schematic outlining subclustering of early developmental lineage populations, including NPC, early neuron, and ChP clusters. **g**, Feature plots displaying YAP1 expression in early developmental lineage populations in control and BD hCOs. **h**, Developmental trajectory mapping of early developmental lineage populations in control and BD hCOs. **i**, Graphical summary of the role of the Hippo signaling pathway in diverting NPCs toward a ChP fate in BD.

To examine whether Hippo pathway activation was restricted to ChP or also present in earlier progenitors, we performed module scoring on the full scRNA-seq dataset. The Hippo module was enriched across the entire clusters, including NPC clusters positioned upstream of ChP differentiation **(Fig. 4b)**. These results suggest that molecular events associated with the Hippo signaling pathway occur prior to ChP differentiation and actively influence this developmental trajectory. To assess whether this early Hippo pathway activity is dysregulated in BD, we compared the expression of key Hippo signaling effectors, *YAP1* and *TEAD1*, between control and BD hCOs. Both genes were markedly upregulated in BD hCOs, particularly within the NPC clusters **(Fig. 4c)**. Guided by this observation, we performed KEGG enrichment analysis specifically within this NPC cluster to identify dysregulated pathways.

Among all pathways analyzed, Hippo signaling pathway emerged as one of key upregulated pathways in BD hCOs **(Fig. 4d)**, along with robust induction of its canonical genes **(Fig. 4e)**. Taken together, these findings indicate that the Hippo signaling pathway is aberrantly activated in BD hCOs as early as the neural progenitor stage, preceding lineage commitment.

To pinpoint the earliest molecular fate determinants for ChP or neural progenitors, we reanalyzed early developmental lineages excluding mature neuron clusters **(Fig. 4f)**. We asked whether Hippo signaling contributes to early cell fate specification by examining the expression dynamics of its key effector, *YAP1*, within these early lineage clusters. Notably, in BD hCOs, *YAP1* expression aligned with the ChP trajectory but was absent from clusters committed to a neuronal fate **(Fig. 4g,h)**, underscoring a selective push toward ChP identity.

Taken together, these sequential analyses define a causal trajectory whereby aberrant Hippo-YAP1 hyperactivation in NPCs shifts the fate decision threshold, promoting excessive commitment to the ChP lineage and accounting for the enlarged ChP fraction observed in BD organoids **(Fig. 4i)**.

### Recurrent Variants in Core Hippo pathway Regulators across Independent BD Cohorts

The consistent dysregulation of the Hippo pathway observed across all BD organoids suggests that this alteration arise from an underlying genetic predisposition rather than from a random differentiation event. To explore this possibility, we investigated genetic variants that may regulate Hippo pathway activity. In a large-scale GWAS analysis (*n* = 41,917 BD; *n* = 371,549 controls)^8^, variants in Hippo pathway regulators, including *CUL4A*, *STK4*, and *MARK2*, were significantly enriched (*P* < 1 × 10^⁻5^). Additionally, a significant variant burden was observed in key effectors such as *YAP1* and *TEAD1* **(Fig. 5a and Extended Data Fig. 8a)**.

**Fig. 5:**
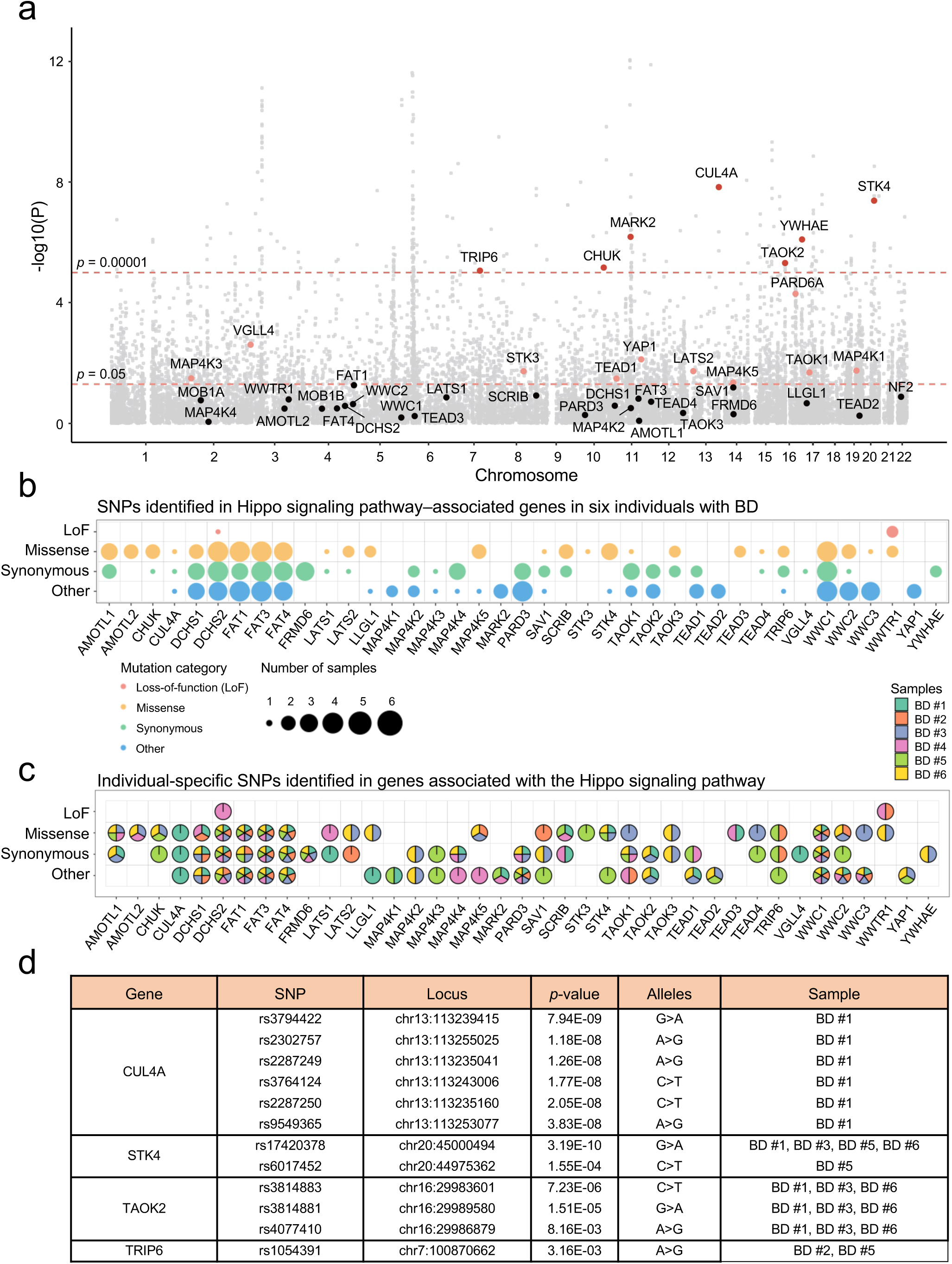
WGS identifies recurrent variants in Hippo pathway genes. **a**, Gene-based Manhattan plot summarizing genetic associations from GWAS in a large BD cohort (*n* = 371,549 controls; *n* = 41,917 BD), with genes annotated to the Hippo signaling pathway specifically highlighted. Red and pink dots represent genes with *P* < 1 × 10⁻⁵ and *P* < 0.05, respectively. **b**, Dot plot providing a summary of SNPs found in Hippo signaling genes across six individuals with BD. Red dots indicate loss-of-function (LoF) variants, yellow dots indicate missense variants, green dots indicate synonymous variants, and blue dots indicate other types of variants. Circle size reflects the number of samples carrying each variant. **c**, Dot plot showing individual-specific SNPs identified in genes associated with the Hippo signaling pathway. Each individuals is represented by a unique color: BD #1 (emerald), BD #2 (coral), BD #3 (light blue), BD #4 (plum), BD #5 (celery), and BD #6 (yellow). **d**, Table summarizing high-priority SNPs from a large BD GWAS cohort that are also identified in six individuals with BD.

To assess the reproducibility of these signals in our samples, we generated WGS data from the six individuals with BD. Variant counts and class distributions (loss-of-function, missense, synonymous, and other) were consistent across all individuals with BD, indicating high and comparable data quality **(Extended data Fig. 8b,c)**. Comparison between our variant catalogue and the public Hippo pathway variant list revealed that most high-confidence variants were present in at least one individual from our cohort, highlighting substantial overlap between population-level genetic signals and those found in our cohort **(Fig. 5b,c)**.

Prioritization of variants with the strongest associations in a public dataset (*P* < 1 × 10^⁻5^) identified *CUL4A*, *STK4*, *TAOK2*, and *TRIP6* as key candidates. A SNP-based GWAS on the same dataset further pinpointed individual SNPs within several Hippo pathway genes **(Extended Data Fig. 8d)**. To assess whether these sentinel SNPs were replicated in our samples, we performed a cross-dataset comparison. Among the variants identified, several overlapped with loci prioritized in public datasets. Notably, the damaging missense variant rs17420378 in *STK4*, a core kinase of the Hippo cascade, was found in the heterozygous state in four of six individuals with BD **(Fig. 5d)**. The recurrence of this variant across independent individuals with BD, coupled with its central position within the signaling cascade, provides genetic support for dysregulation of the Hippo signaling pathway.

Altogether, these findings offer compelling molecular evidence that inherited genetic “priming” reduces the activation threshold of the Hippo pathway in NPCs. This shift biases developmental trajectories toward the ChP lineage with Hippo hyperactivation as a causal, rather than correlative, driver of disease pathogenesis.

### Disrupted Cell Junctions in BD hChPOs

Previous studies have reported disrupted cell junctions in individuals with BD, implicating impaired epithelial barrier function^19^. Given the importance of junctional integrity for ChP function and the role of Hippo signaling in epithelial architecture^20^, we examined junction-related gene expression in BD hChPOs. RNA-seq analysis revealed dysregulation of cell junction genes, including the tight junction component *CLDN2* **(Extended Data Fig. 9a,b)**. Gene Ontology (GO) analysis revealed significant enrichment of terms related to cell junctions, including cell-cell junction organization and cell junction assembly **(Fig. 6a)**. Notably, many DEGs were associated with cell junctions, most of which were significantly downregulated in BD hChPOs **(Fig. 6b)**. Core structural components of multiple junction types, including *CTNNB1* (adherens junction), *TJP1* (tight junction), *DSP* (desmosome), and *VCL* (gap junction), were markedly downregulated in BD hChPOs **(Fig. 6c)**. KEGG pathway analysis similarly showed decreased expression in tight junction and gap junction pathways in BD hChPOs **(Fig. 6d)**.

**Fig. 6:**
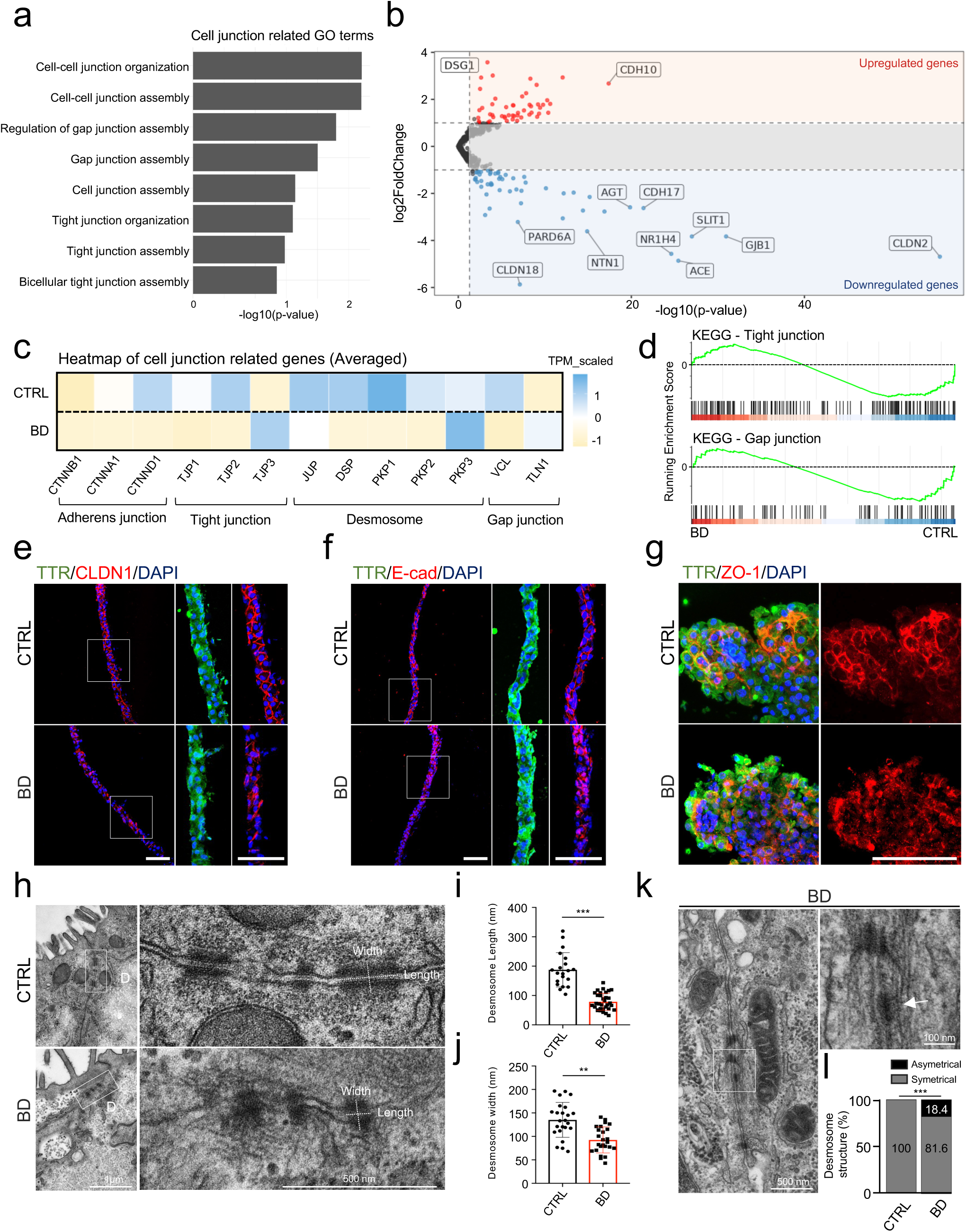
BD hChPOs exhibit disrupted cell junctions. **a**, GO terms associated with cell junction–related processes. **b**, Differential expression of cell junction–related genes in BD hChPOs compared to control hChPOs. Red dots indicate upregulated genes (log_2_ fold change > 1, FDR < 0.05), and blue dots indicate downregulated genes (log_2_ fold change < -1, FDR < 0.05) in BD hChPOs. **c**, Heatmap showing expression levels of cell junction–related genes. **d**, GSEA of KEGG pathways related to tight junction and gap junction in control and BD hChPOs. **e,f**, Immunofluorescence images of control and BD hChPOs at Day 80 stained for TTR in combination with CLDN1 (e) and E-cad (f). Nuclei are counterstained with DAPI. The white box indicates the magnified region shown on the right. Scale bars, 100 µm. **g**, Whole-mount immunofluorescence images of control and BD hChPOs stained for TTR and ZO-1. Nuclei are counterstained with DAPI. Scale bars, 100 µm. **h**, Ultrastructural images of cell junctions in control and BD hChPOs. The white box indicates the magnified region shown on the right. High-magnification images of desmosomes are shown on the right. Scale bars, 1 µm (left), 500 nm (right). **i**, Quantification of desmosome length in control (*n* = 20) and BD (*n* = 33) hChPOs. **j**, Quantification of desmosome width in control (*n* = 24) and BD (*n* = 25) hChPOs. **k**, Ultrastructural images of desmosomes in BD hChPOs. The white box indicates the magnified region shown in the upper right. The white arrow highlights the asymmetric structure of the desmosome in the high-magnification image. Scale bars, 500 nm (left), 100 nm (upper right). **l**, Quantification of desmosome structural distributions in control (*n* = 30) and BD (*n* = 38) hChPOs. All data are presented as mean ± s.d. Statistical analysis was performed using a two-tailed Student’s t-test (i,j,l). \*\*\**P* < 0.001, \*\**P* < 0.01. D, desmosome.

Given these transcriptional alterations, we next examined the structural integrity of cell junctions in BD hChPOs to determine whether these changes are reflected in cellular architecture. Notably, CLDN1 expression exhibited a disrupted and disorganized pattern in all BD hChPOs, in contrast to the continuous junctional staining observed along cell-cell borders in control hChPOs **(Fig. 6e and Extended Data Fig. 9c)**. In control hChPOs, E-cadherin and ZO-1 were clearly localized at cell-cell interfaces, forming continuous adherens and tight junctions. In BD hChPOs, however, staining for E-cadherin and ZO-1 appeared irregular and discontinuous, with reduced membrane localization, further supporting the presence of disrupted junctional architecture in BD **(Fig. 6f,g)**.

To further elucidate the structural abnormalities in cell junctions, we examined their ultrastructural organization. In control hChPOs, cell-cell junctions displayed a canonical apical-to-basal organization, with clearly distinguishable tight junctions, adherens junctions, and desmosomes. In contrast, BD hChPOs exhibited pronounced structural abnormalities, including poorly defined tight and adherens junctions, and desmosomes that frequently appeared clustered rather than uniformly distributed **(Extended Data Fig. 9d)**. Among these abnormalities, desmosomes were particularly affected in BD hChPOs, with individual structures showing marked reductions in both length and width **(Fig. 6h)**. Quantitative analysis confirmed that desmosomes in BD hChPOs were significantly shorter and narrower compared to control hChPOs **(Fig. 6i,j)**. Additionally, BD hChPOs exhibited severely disrupted desmosome architecture, often presenting as asymmetric structures that lacked one side of either the intercellular plaques or the intermediate filament connection **(Fig. 6k and Extended Data Fig. 9e,f)**. Quantitative analysis further demonstrated a significant increase in the proportion of asymmetric desmosome structures in BD hChPOs **(Fig. 6l and Extended Data Fig. 9g)**.

Collectively, these data reveal that cell junction architecture is markedly compromised in BD hChPOs, a defect that underline or exacerbate ChP dysfunction in individuals with BD.

### Altered iCSF Composition in BD hChPOs

The CSF secreted by the ChP plays a vital role in maintaining brain homeostasis through nutrient exchange, waste clearance, and intercellular signaling^21^. We therefore examined whether ChP alterations in BD contribute to the CSF dysfunction and compositional differences. To this end, we collected iCSF secreted into the lumens of control and BD hChPOs and performed proteomic profiling at Day 80 **(Fig. 7a)**. To assess the biological relevance of the iCSF, we first compared its protein composition to previously published hChPO-derived iCSF^18^ and *in vivo* human CSF proteomic datasets^22–24^. A total of 807 proteins were identified in hChPO-derived iCSF, filtered by high-confidence criteria (≥2 unique peptides and false discovery rate (FDR) ≤1%) **(Supplementary Table 2)**. Consistent with a previous study^18^, GO analysis revealed functional enrichment in multiple categories, including cell adhesion molecule binding (MF), extracellular vesicle (CC), and key signaling and metabolic pathways such as carbon metabolism (KEGG) and VEGFA–VEGFR2 signaling (WP) **(Fig. 7b)**. Moreover, a significant proportion of proteins overlapped with published hChPO-derived iCSF and *in vivo* human CSF datasets **(Fig. 7c)**, highlighting the biological fidelity of our iCSF model. Notably, most identified proteins (490 out of 807) absent from the culture medium **(Fig. 7d, Extended Data Fig. 10a and Supplementary Table 3)**, indicating de novo secretion by hChPOs, for example, SERPINF1, IGF2, and LDHB **(Extended Data Fig. 10b)**. In addition, substantial overlap with normal human embryonic, pediatric, and adult CSF, particularly adult CSF, reinforces the physiological relevance of our iCSF model **(Fig. 7e, Extended Data Fig. 10c and Supplementary Table 4)**.

**Fig. 7:**
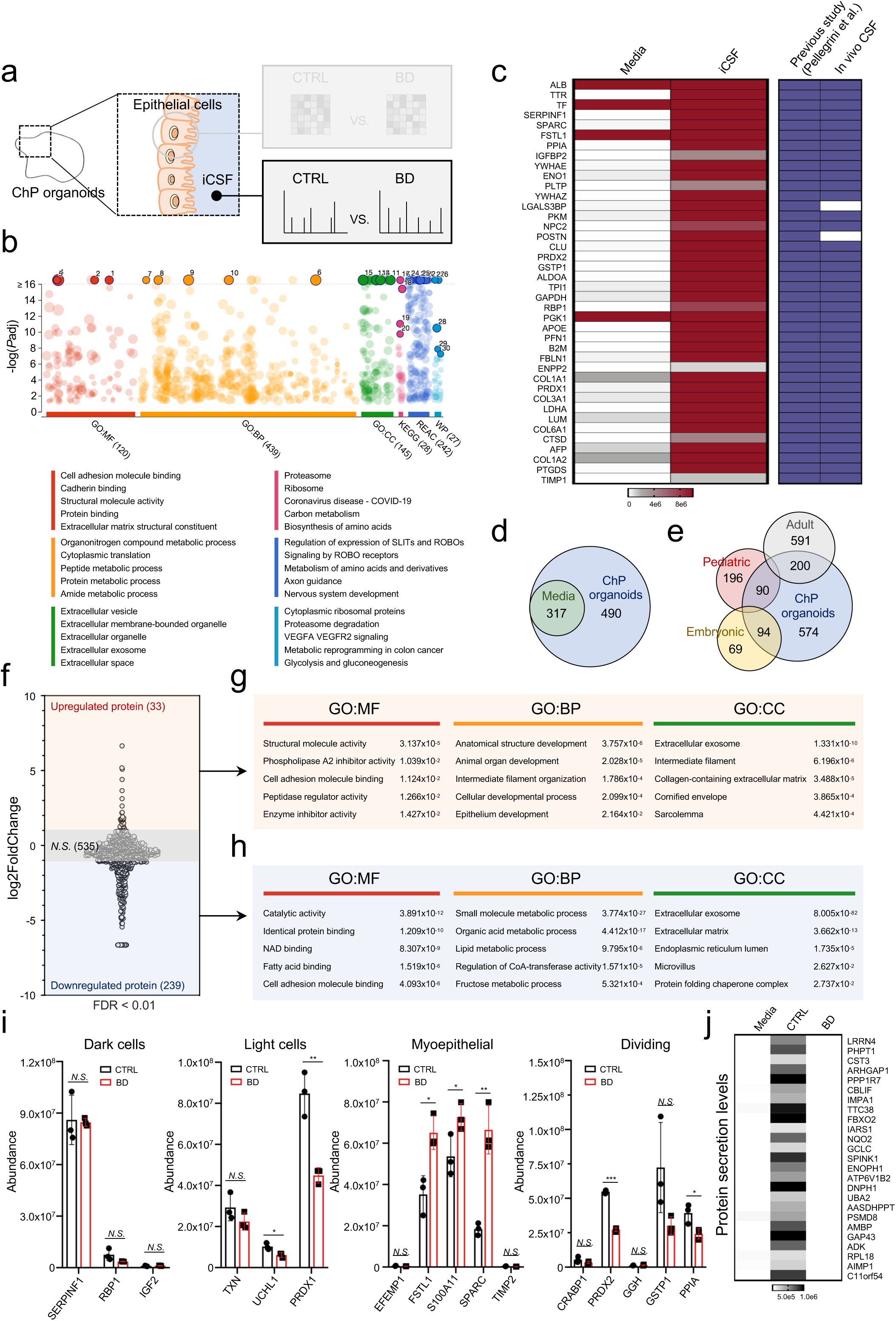
BD hChPOs exhibit altered iCSF composition. **a**, Schematic outlining the proteomic analysis of control and BD hChPOs at Day 80. **b**, g:ProfileR analysis of proteins identified in iCSF. The top five enriched terms in each category are highlighted, with categories distinguished by color: red for molecular function (MF), yellow for biological process (BP), green for cellular component (CC), magenta for KEGG, blue for Reactome (REAC), and turquoise for WikiPathways (WP). **c**, Heatmap showing representative proteins identified in iCSF (red), compared to previously published iCSF and in vivo CSF datasets (purple). **d**, Venn diagram illustrating shared (green) and uniquely identified proteins (blue) between iCSF and media. **e**, Venn diagram showing the overlap between proteins identified in iCSF (blue) and proteomic datasets from human adult (grey), pediatric (red), and embryonic (yellow) CSF. **f**, DAPs in BD iCSF compared to control iCSF. Upregulated proteins are highlighted in red-shaded areas (log_2_ fold change > 1, FDR < 0.01), and downregulated proteins are highlighted in blue-shaded areas (log_2_ fold change < -1, FDR < 0.01). Proteins within the grey-shaded area are not significantly different (*N.S.*). **g**,**h**, GO enrichment analysis of upregulated proteins (g) and downregulated proteins (h) in BD iCSF. The five representative enriched terms from each category are shown. **i**, Comparative abundance of ChP epithelial cell type-specific proteins in control and BD iCSF (*n* = 3 per group). **j**, Heatmap showing specific proteins undetected in BD iCSF. All data are presented as mean ± s.d. Statistical analysis was performed using a two-tailed Student’s t-test (i). \*\*\**P* < 0.001, \*\**P* < 0.01, \**P* < 0.05, *N.S*., not significant.

We next investigated BD-associated changes by comparing BD and control iCSF proteomes, identifying 272 differentially abundant proteins (DAPs), including 33 upregulated and 239 downregulated proteins in BD iCSF **(Fig. 7f, Supplementary Tables 5 and 6)**. Notably, 73 DAPs overlapped with developmental stage-specific CSF proteins, underscoring the utility of hChPOs in exploring the temporal dynamics of BD-associated CSF proteins **(Extended Data Fig. 10d)**. GO analysis revealed that upregulated proteins in BD iCSF were enriched in developmental processes, including anatomical structure development, cellular developmental process, and epithelium development **(Fig. 7g)**, reflecting abnormal ChP development. Downregulated proteins were significantly underrepresented in metabolic processes such as small molecule metabolic process, lipid metabolic process, and fructose metabolic process **(Fig. 7h)**, suggesting impaired metabolic activity. Together, these alterations in the iCSF proteome may influence neuronal function by perturbing developmental and metabolic signaling pathways relevant to BD.

A previous study identified four distinct epithelial cell types within hChPOs that contribute to CSF secretion^18^. To investigate cell-type-specific alterations in BD, we analyzed changes in cell type-specific protein secretion. Dark cell-specific proteins such as SERPINF1 and RBP1 were consistently secreted in both control and BD hChPOs, whereas the secretion levels of light cell-specific proteins, including UCHL1 and PRDX1, were reduced in BD hChPOs. In addition, the secretion levels of myoepithelial cell-specific proteins, such as FSTL1 and SPARC, were increased, while proteins associated with dividing cells, including PRDX2 and GSTP1, showed decreased secretion in BD hChPOs **(Fig. 7i and Supplementary Table 7)**. Importantly, these secretion profiles were closely mirrored with gene expression changes at the RNA level **(Extended Data Fig. 10e)**. These findings suggest that cell type-specific protein secretion is altered in BD hChPOs.

Consistent with a previous study^18^, our iCSF dataset also included clinically relevant biomarkers such as APOE (Alzheimer’s disease), PARK7 (Parkinson’s disease), and HSPA5 (glioma) **(Extended Data Fig. 10f and Supplementary Table 8)**, prompting further examination of BD-associated protein alterations in iCSF. Notably, several BD-associated proteins, including APOA1 and A2M^25,26^, exhibited significant differences in BD iCSF **(Extended Data Fig. 10g and Supplementary Table 9)**, suggesting their relevance to BD-related molecular pathology. Furthermore, we observed 25 proteins that were undetectable in BD iCSF, including the neuron-specific growth-associated protein GAP43 **(Fig. 7j and Supplementary Table 10)**, suggesting a possible link to BD pathology.

Altogether, these data suggest that BD is associated with changes in the iCSF proteome that include altered cell type-specific protein secretion and dysregulation of neurodevelopmental and metabolic pathways, which may contribute to BD pathology.

## Discussion

Our study reveals a previously unappreciated early neurodevelopmental bias toward a ChP fate in BD. Through single-cell transcriptomics, we show that BD hCOs consistently exhibit ChP lineage expansion. This expansion is validated in an independent hChPO model. The finding is also corroborated by *in vivo* neuroimaging data from the same individuals. Mechanistically, ChP overgrowth is driven by hyperactivation of the Hippo signaling pathway representing a convergent mechanism. Trajectory analysis reveals that this dysregulation arises in NPCs prior to lineage bifurcation, suggesting that these cells are primed toward ChP identity. Notably, WGS revealed recurrent genetic variants in Hippo regulators, including *STK4* and *YAP1*, across all six individuals with BD. These genetic variants may contribute to Hippo pathway dysregulation, thereby providing a potential genetic basis for the observed ChP expansion in BD.

An integrative multi-omics approach reveals functional consequences of these molecular alterations in BD hChPOs, which exhibit disrupted epithelial junctions and aberrant iCSF secretion. Compromised epithelial integrity, a hallmark of ChP dysfunction, was confirmed at the ultrastructural level. Proteomic profiling of iCSF revealed altered secretion patterns, including cell type-specific changes and dysregulation of developmental and metabolic pathways. Notably, 25 proteins were absent in BD iCSF, most not previously implicated in BD. Among these, the neuron-specific growth-associated protein GAP43—previously reported as reduced in BD postmortem brains^27^—and the cysteine protease inhibitor CST3, which has been reported to be decreased in the CSF of patients with depression^28^, were undetectable, consistent with impaired synaptic plasticity and altered CSF proteostasis. Overall, these findings highlight not only the pathological relevance of iCSF alterations in BD but also underscore the potential of hChPO-derived iCSF as a translational platform for biomarker discovery.

Taken together, these findings converge on a model in which inherited vulnerability in Hippo signaling primes early NPCs toward a ChP fate, resulting in structural and functional aberrations of the ChP that manifest in both organoid models and clinical imaging. This work provides compelling evidence that the ChP, historically overlooked in BD research, plays an active role in disease pathogenesis and may serve as a novel biomarker source and therapeutic target. By leveraging, patient-derived iPSCs and standardized organoid differentiation protocols, our platform minimizes environmental variability and isolates the contribution of inherited genetic factors. This controlled system enables direct interrogation of genotype-driven mechanisms underlying ChP fate bias and Hippo pathway dysregulation in BD.

Nevertheless, several limitations should be noted. The sample size in our study was modest; however, the convergence of findings across multiple complementary datasets, including genomic, transcriptomic, and imaging analyses, supports the robustness of our conclusions. The sample was limited to females so it is not clear if findings will extend to males. Furthermore, while the iCSF proteome recapitulates many features of native CSF, direct validation with patient-derived CSF samples would further strengthen the translational relevance of our model. Finally, the downstream impact of altered ChP function on neuronal circuits and behavior remains to be elucidated.

In conclusion, we present a comprehensive, multimodal investigation that implicates the ChP as an active and genetically primed contributor to the early pathogenesis of BD. By integrating genomic, transcriptomic, proteomic, ultrastructural, and imaging data, our study uncovers a robust ChP fate bias in BD, driven by Hippo pathway hyperactivation and genetic variants in its key regulators, thereby culminating in both structural disorganization and functional impairment of the ChP. These findings expand on neurocentric models of BD by positioning the ChP as a critical driver of BD brain pathology. Our work not only broadens the conceptual framework for BD pathophysiology but also establishes patient-derived organoids as a versatile translational platform for advancing both diagnostic and therapeutic strategies.

## Methods

### Human participants and *in vivo* magnetic resonance imaging

Structural magnetic resonance images (sMRI) were acquired from six subjects with BD (mean age 54.8±11.7 years) and three control subjects (52.3±4.3 years). All subjects were female. The presence or absence of Axis I diagnoses was confirmed with the Structured Clinical Interview (SCID) for Diagnostic and Statistical Manual of Mental Disorders, Fifth Edition (DSM-5) – Research Version^29^. The control subjects included participants without a lifetime history of Axis-I disorders or first-degree relatives with a major psychiatric disorder, as assessed using the Family History Screen for Epidemiological Studies^30^. Exclusion criteria for all participants were major medical or neurologic disorders that could affect brain tissue. Participants provided written informed consent in accordance with the Yale School of Medicine Human Investigation Committee/Institutional Review Board.

For the BD participants, all had mood symptoms by the adolescent/early adult epoch when fully syndromic BD often first emerges^31^ and a first-degree relative with BD (range 1–3 relatives). At scanning, three BD participants were euthymic, one was depressed and one was manic. Three BD participants had a history of psychotic symptoms during at least one mood episode^32^ and one had a history of rapid-cycling. Four BD participants had a history of at least one psychiatric hospitalization (range 1–5). Three had a history of suicide attempt, determined using the Columbia Suicide History Form^33^. Five BD participants were taking psychotropic medications that included antidepressants (*n* = 4), anticonvulsants (*n* = 3), a second-generation antipsychotic (*n* = 3), a benzodiazepine (*n* = 3), and an opiate (*n* = 1). One BD subject had hypothyroidism, which is common in BD, and was treated with levothyroxine.

MRI data were acquired with a high-resolution 3D T1-weighted magnetization-prepared rapid gradient echo (MPRAGE) sequence using a 3-Tesla Prisma MRI Scanner (Siemens, Erlangen, Germany) with parameters: repetition time (TR)=2500 ms, echo time (TE)=2.81 ms, matrix=256×256, slices=176, voxel size=1.0×1.0×1.0 mm^3^, no intersection gap, slice thickness=1.00 mm, flip angle (FA)=7°, field of view (FOV)=256×256 mm^2^, and scan time=8.02 min. Preprocessing was conducted using FreeSurfer v6.0.0 (https://surfer.nmr.mgh.harvard.edu) via the autorecon pipeline, which includes motion correction, intensity normalization, Talairach transformation, skull stripping, gray/white matter boundary tessellation, subcortical white matter segmentation, automated topology correction, and cortical surface reconstruction^34,35^. LV volumes and ChP segmentation were quantified using Gaussian Mixture Models (GMM)^36^. GMM, an unsupervised machine learning approach^37^, was applied to all voxels within the LVs to distinguish CSF, ventricular wall, and ChP tissue. The modeling was initiated using a mask that encompassed all relevant voxels, created by merging ventricular and ChP segmentations generated with FreeSurfer. A Bayesian GMM with two components (implemented using the scikit-learn Python library)^38^ was then applied to all voxels within the mask. This procedure classified voxels into two groups: one with lower mean intensity values, primarily representing CSF, and another with higher intensities, corresponding to ChP and ventricular wall regions. Voxels assigned to the higher-intensity cluster were subsequently smoothed using the 3D Smallest Univalue Segment Assimilating Nucleus (SUSAN) algorithm from FSL (σ=1 mm)^39^. This step leverages spatial context: ventricular wall voxels are adjacent to CSF (assigned intensity=0), leading to a reduction in intensity after smoothing, whereas ChP voxels are surrounded by similar tissue (intensity=1), preserving higher intensities. Finally, a second Bayesian GMM with three components was applied to the smoothed voxels, and those within the highest-intensity cluster were identified as ChP tissue. For analyses, all volume values were the percentage with respect to the total intracranial volume^40^.

### iPSC generation and characterization

Human iPSCs were generated from peripheral blood mononuclear cells (PBMCs) using the CytoTune-iPS 2.0 Sendai Reprogramming Kit (A16517, Thermo Fisher Scientific), which utilizes a non-integrating Sendai virus–based system, according to the manufacturer’s instructions. Briefly, cells were transduced with Sendai vectors encoding *OCT4*, *SOX2*, *KLF4*, and *c-MYC* and cultured on Matrigel-coated plates in mTeSR1 medium (STEMCELL Technologies). Colonies exhibiting human embryonic stem cell (hESC)-like morphology emerged between days 18 and 25 post-transduction and were manually picked and expanded. Once established, all iPSCs were maintained in mTeSR1 medium with daily medium changes. Pluripotency of all established iPSC lines was assessed by immunostaining and RT-qPCR for key pluripotency markers, including OCT4 and NANOG. All procedures were conducted in accordance with approved safety protocols of the Yale School of Medicine.

### RNA extraction and RT-qPCR

Total RNA was extracted using the RNeasy Mini Kit (Qiagen), and 1 µg of RNA was reverse transcribed into cDNA with the ReverTra Ace™ qPCR RT Master Mix (FSQ-201, TOYOBO). Quantitative real-time PCR was performed using the iTaq Universal SYBR Green Supermix (Bio-Rad) on a CFX384 Real-Time PCR System (Bio-Rad). The thermal cycling protocol consisted of an initial denaturation at 95°C for 15 min, followed by 40 cycles of 94°C for 10 sec and 60°C for 45 sec. Primer sequences used for gene expression analysis are provided below: OCT4 (forward primer: 5’-GACAGGGGGAGGGGAGGAGCTAGG-3’; reverse primer: 5’-CTTCCCTCCAACCAGTTGCCCCAAAC-3’), NANOG (forward primer: 5’-CATGCAACCTGAAGACGTGTGA-3’; reverse primer: 5’-AATCCTATGAGGGATGGGAGGA-3’), SOX2 (forward primer: 5’-GGGAAATGGGAGGGGTGCAAAAGAGG-3’, reverse primer: 5’-TTGCGTGAGTGTGGATGGGATTGGTG-3’), and REX1 (forward primer: 5’-CTGAGGCTGGAGCCTGTGTG-3’, reverse primer: 5’-AACCACCTCCAGGCAGTAGTGA-3’).

### Brain organoid generation

For hCO generation, iPSCs were dissociated using Accutase and counted with the ADAM-CellT system (NanoEntek). A total of 9,000 cells per well were seeded into low-attachment, U-bottom 96-well plates (Corning) containing neural induction medium composed of DMEM/F12, 15% (v/v) KSR, 5% (v/v) heat-inactivated FBS (Life Technologies), 1% (v/v) GlutaMAX, 1% (v/v) MEM-NEAA, and 100 µM β-mercaptoethanol, supplemented with 10 µM SB431542, 100 nM LDN193189, 2 µM XAV939, and 50 µM Y-27632. FBS was removed from the medium on day 2, and Y-27632 was withdrawn on day 4. Media were refreshed every other day until day 10, at which point the organoids were transferred to ultra-low-attachment six-well plates and cultured on orbital shakers in cortical patterning medium. This medium consisted of a 1:1 mixture of DMEM/F12 and Neurobasal medium supplemented with 0.5% (v/v) N2 supplement, 1% (v/v) B27 supplement without vitamin A, 0.5% (v/v) MEM-NEAA, 1% (v/v) GlutaMAX, 50 µM β-mercaptoethanol, 1% (v/v) penicillin/streptomycin, and 0.025% insulin. From day 18 onward, the medium was replaced with cortical differentiation medium containing B27 supplement with vitamin A and further supplemented with 20 ng/mL BDNF and 200 µM ascorbic acid. Media were changed every 3 days.

For hChPO generation, organoids were generated from all iPSC lines following a previously established protocol^18^, with minor modifications. A total of 9,000 cells per well were seeded into low-attachment, U-bottom 96-well plates containing embryoid body (EB) media supplemented with 50 μM Y-27632 for the first 3 days, followed by 2 days in EB media alone. EB media consisted of DMEM/F12 supplemented with 20% KSR, 1% GlutaMAX, 1% MEM-NEAA, and 0.1 mM β-mercaptoethanol. On day 5, EBs were transferred to neural induction media composed of DMEM/F12, 1% N2 supplement, 1% GlutaMAX, 1% MEM-NEAA, and 1 μg/ml heparin. After two days, EBs were embedded in Matrigel and transferred to expansion media consisting of a 1:1 mixture of DMEM/F12 and Neurobasal medium, supplemented with 0.5% (v/v) N2 supplement, 1% (v/v) B27 supplement without vitamin A, 0.5% (v/v) MEM-NEAA, 1% (v/v) GlutaMAX, 50 µM β-mercaptoethanol, 1% (v/v) penicillin/streptomycin, and 0.025% insulin. For ChP patterning, organoids were treated on day 10 with 3 μM CHIR99021 and 20 ng/mL BMP4 for 7 days. From day 10 onward, maturation medium was used, identical to the expansion medium except that B27 supplement with vitamin A was substituted. Organoids were transferred to orbital shakers beginning on day 15. From day 30, dissolved Matrigel was added to the maturation media. Media were changed every 3 days.

### Immunohistochemistry and confocal imaging

Whole organoids were fixed in 4% paraformaldehyde (PFA) at 4°C overnight and washed three times with PBS. Samples were cryoprotected in 30% sucrose at 4°C for 48 h, embedded in optimal cutting temperature (OCT) compound, and cryosectioned at 30-40 μm thickness using a cryostat. Sections were mounted on glass slides and permeabilized with 0.1% Triton X-100 in PBS, followed by blocking with 3% bovine serum albumin (BSA) for 2 h at room temperature. Primary antibodies were applied and incubated overnight at 4°C. After PBS washes, Alexa Fluor–conjugated secondary antibodies (1:1000) were applied for 2 h at room temperature. Nuclei were counterstained with DAPI (1:1000) for 10 min, and sections were mounted using VECTASHIELD (Vector Laboratories). Imaging was performed using a STELLARIS WLL confocal microscope (Leica Microsystems). The following primary antibodies were used in this study: Goat anti-SOX2 (AF2018, R&D Systems), Mouse anti-TTR (MAB7505, R&D Systems), Rabbit anti-LMX1A (PA5-115517, Invitrogen), Rabbit anti-Claudin1 (ab211737, Abcam), Rabbit anti-E-Cadherin (ab40772, Abcam), Rabbit anti-ZO1 (ab221547, Abcam), Rabbit anti-active YAP1 (ab205270, Abcam).

### Whole-mount immunofluorescence staining

Whole organoids were fixed in 4% PFA for 1 h, washed with PBS, and permeabilized in 0.1% Triton X-100 overnight at 4°C. After blocking with 3% BSA, samples were incubated with primary antibodies for 48 h at 4°C, followed by overnight incubation with appropriate secondary antibodies (1:500). Nuclei were counterstained with DAPI (1:500) for 1 h. Prior to imaging, samples were immersed in a clearing solution (25% urea and 65% sucrose in water) to match the refractive index (RI).

### Electron Microscopy (EM) Imaging

Organoids were fixed in a solution containing 2.5% glutaraldehyde, 2.0% PFA in 0.1 M sodium cacodylate buffer (pH 7.4) followed by rinsing in the same buffer. Post-fixation was performed in a solution of 1% osmium tetroxide (Electron Microscopy Sciences, EMS) and 0.8% potassium ferrocyanide in 0.1 M sodium cacodylate buffer (pH 7.4). After thorough rinsing in 0.1 M sodium cacodylate and HPLC-grade water, samples were stained en bloc with 2% uranyl acetate (EMS) and subsequently rinsed in HPLC water. Samples were dehydrated through a graded ethanol series (50%, 70%, 90%, and 100%), transitioned to propylene oxide (EMS), and infiltrated overnight in EPON Embed 812 epoxy resin (EMS). When the organoids were fully infiltrated with EPON, resin was polymerized at 60°C overnight. Ultrathin sections (60 nm) were prepared using a Leica UC7 ultramicrotome, collected on formvar-coated copper slot grids, and post-stained with 2% uranyl acetate and Reynolds’ lead citrate (EMS).

Samples were imaged on a Tecnai G2 Spirit BioTwin Transmission electron microscope (Thermo Fisher Scientific) operating at 80 kV. Images were acquired using a NanoSprint15 MKII camera (AMT Imaging) and AMT Capture Engine Software (v7.0.2.5).

### Library preparation and sequencing for WGS analysis

WGS was performed on genomic DNA from six individuals with BD using the Yale Center for Genome Analysis (YCGA) pipeline. Following standard assessments of DNA quantity, quality, and purity, samples were further quantified using the Qubit dsDNA assay (Q33226, Thermo Fisher Scientific) to ensure compatibility with PCR-free library preparation protocols. Library preparation was performed using 250 ng of high-quality genomic DNA with the Watchmaker DNA Library Prep Kit (7K0101-096, Watchmaker Genomics), which enables enzymatic fragmentation, end-repair, and A-tailing in a single reaction. xGen UDI-UMI Adapters (10005903, IDT) were ligated to the fragmented DNA for dual indexing. Final library construct was assessed using the Caliper LabChip GX system, and library quantification was performed by qPCR with the Kapa Library Quantification Kit (KK4854, Roche). Normalized libraries (2 nM) were sequenced on an Illumina NovaSeq X Plus system using 151 bp paired-end reads. Libraries were sequenced to a depth of at least 100 Gbp per sample using an optimized loading concentration to minimize duplication rates. Dual 10 bp indices were read during dedicated index reads. A 1% PhiX spike-in control was included in each lane to monitor sequencing quality. Signal intensities were processed into base calls using Illumina’s Real-Time Analysis (RTA) software, and subsequent demultiplexing and alignment to the human genome were performed using CASAVA v1.8.2.

### WGS data analysis

Sequencing reads were aligned to the GRCh38 human reference genome using BWA-MEM. Duplicate reads were marked with Picard, and base quality score recalibration was performed using the Genome Analysis Toolkit (GATK), following the GATK Best Practices. SNVs and small indels were called using GATK HaplotypeCaller in gVCF mode, followed by joint genotyping across all samples. Variant annotation was conducted with the Ensembl Variant Effect Predictor (VEP v85).

To ensure a high-confidence callset, variants were subjected to rigorous post-calling quality control. Homozygous alternate genotypes were retained if they passed variant quality score recalibration (VQSR), exhibited an alternate allele depth (AD) ≥ 3, a total read depth (DP) ≥ 10, genotype quality (GQ) ≥ 20, and an allelic balance (AB) ≥ 0.8. For heterozygous genotypes, the same filters were applied, with a relaxed allelic balance threshold (AB ≥ 0.2).

### GWAS analysis

Summary statistics for BD were obtained from the Psychiatric Genomics Consortium (PGC), based on a GWAS comprising 41,917 cases and 371,549 controls of European ancestry^8^. The dataset is publicly available through the PGC data portal (https://pgc.unc.edu/for-researchers/download-results/). Gene-based association analysis was conducted using MAGMA (v1.08)^41^, as implemented in FUMA^42^, with default settings and a zero window size around each gene. Manhattan plots were generated using the ggplot2 package in R (version 4.2.0)^43^.

### scRNA-seq library preparation and sequencing

For four control samples (control #1-4) and six BD samples (BD #1-6), hCOs were randomly collected from independent culture dishes, and five organoids were pooled per sample for dissociation at Day 75. Organoid dissociation, cDNA preparation, and sequencing were performed following previously established protocols. Briefly, organoids were rinsed with HBSS, minced into small fragments, and enzymatically dissociated using papain (Worthington Biochemical Corporation) according to the manufacturer’s instructions. All solutions were pre-oxygenated with 95% O₂ and 5% CO₂ for 5 min. Tissue fragments were further oxygenated for an additional 5 min, then incubated in oxygenated papain solution at 37°C in a water bath for 1 h with gentle agitation every 10 min. The enzymatic reaction was quenched with an albumin-ovomucoid inhibitor solution, and the resulting single-cell suspension was prepared in 1% BSA/PBS containing 10 µM Y-27632. Cells were stained with propidium iodide (PI) for 15 min on ice, sorted to remove dead cells, and resuspended at a final concentration of 1,000 cells/µL. cDNA libraries were prepared using the Chromium Next-GEM Single Cell 3′ v3.1 Reagent Kit (10x Genomics) according to the manufacturer’s protocol. Barcoded, full-length cDNA was synthesized from polyadenylated mRNA, followed by size selection and adaptor ligation, during which R2, P5, and P7 sequences were incorporated. Libraries were amplified using Illumina bridge amplification and sequenced on an Illumina NovaSeq S4 platform (2 × 150 bp, Rapid Run Mode).

### scRNA-seq data analysis

To obtain count matrices, scRNA-seq reads were aligned to GRCh38 human genome and counted with Ensembl genes by count function of CellRanger (v 6.1.2) with default parameters, generating gene-barcode count matrices. Subsequently, the Read10X function of Seurat (v 4.3.0)^44^ was used to import the count matrices for four control samples (control #1-4) and six BD samples (BD #1-6). Quality control was performed to remove low-quality cells. Additionally, cells with an excessively high percentage of mitochondrial gene expression (above 10%) were excluded. Following cell filtering, normalization was performed with the NormalizeData function in Seurat, using a scaling factor of 10,000 and log-transformation based on total cell expression. Next, highly variable genes were identified using the FindVariableFeatures function with the VST method. To account for batch effects across samples, Seurat’s anchor-based integration workflow was employed, including SelectIntegrationFeatures, FindIntegrationAnchors, and IntegrateData. The integrated dataset was then scaled using ScaleData, and principal component analysis (RunPCA) was conducted with 30 principal components. Clustering was performed with FindNeighbors and FindClusters (resolution = 0.7), followed by dimensionality reduction via RunUMAP using the same principal components. As a result of this process, a total of 25 clusters were identified and subsequently annotated according to general organoid markers. First, based on the expression levels of SLC17A6 and SLC17A7, mature cortical (excitatory) neuron clusters were classified. Additionally, InN clusters were defined according to the expression levels of GAD1 and GAD2. General neuron markers such as DCX and MAP2 was confirmed neuronal clusters. NPC clusters were identified based on the expression of the general NPC markers and early proliferation markers PAX6, SOX2, MKI67, and TOP2A. ChP clusters were divided based on TTR, CLIC6, respectively. For trajectory analysis, the Monocle3 (v1.3.7) package was used^45^. First, the expression matrix was extracted from the Seurat object and converted into a sparseMatrix format. The cell metadata and gene annotation were retrieved from the Seurat object, and a cell_data_set (CDS) was created using the new_cell_data_set function. Cells were then clustered, and a trajectory graph was learned using learn_graph. Pseudotime ordering was performed using order_cells with the UMAP reduction method.

### Bulk RNA-seq

Total RNA was extracted from whole organoids at Day 80 using the RNeasy Mini Kit (Qiagen, 74106) following the manufacturer’s protocol. RNA quality and integrity were assessed using the Agilent Bioanalyzer 2100 (Agilent Technologies), and only samples with an RNA Integrity Number (RIN) ≥ 8 were used for library construction. RNA-seq libraries were prepared using the TruSeq Stranded mRNA Library Preparation Kit (Illumina) according to the manufacturer’s instructions, and sequencing was performed on the Illumina HiSeq 4000 platform.

For data preprocessing, paired-end raw reads were trimmed using fastp (v0.23.4) with default parameters. The quality of trimmed reads was evaluated using FastQC (v0.12.1). Reads were then aligned to the GRCh38 human reference genome using STAR aligner (v2.7.11b)^46^, producing both genome-aligned and transcriptome-aligned BAM files. The resulting BAM files were sorted and indexed using samtools (v1.20)^47^. Transcript quantification was performed in alignment-based mode using Salmon (v1.10.3)^48^, with transcript annotations provided via a GFFread-generated reference or pre-existing FASTA file (specified with the -t parameter). All intermediate and final outputs, including trimmed FASTQ files, alignment statistics, sorted/indexed BAMs, and quantified expression values, were organized for downstream processing.

DEG analysis was conducted using DESeq2 (v1.38.3)^49^. Genes with a FDR < 0.05 and a log₂(fold-change) > 1 or < –1 were considered significantly differentially expressed.

### Mass spectrometry (MS) analysis

Proteomic profiling was performed on Day 80 iCSF samples from control and BD hChPOs, with biological triplicates for each group. Samples were dried, denatured in urea, reduced with dithiothreitol (DTT), and alkylated with iodoacetamide to prevent disulfide bond formation. Proteins were digested overnight with Lys-C at 37°C, followed by an additional 4 h digestion with trypsin to generate peptides. Peptides were acidified, desalted using BioPureSPN PROTO 300 C18 MacroSpin columns (The Nest Group), and dried via SpeedVac. Dried peptides were reconstituted in MS loading buffer solution. Peptides were analyzed on a Thermo Scientific Q Exactive HFX mass spectrometer coupled to a Waters nanoAcquity UPLC system using a binary solvent system. Trapping was performed at 5 µL/min on a Waters Symmetry C18 trap column (100 Å, 5 µm, 180 µm × 20 mm). Analytical separation was carried out at 37°C on an ACQUITY UPLC Peptide BEH C18 column (130 Å, 1.7 µm, 75 µm × 250 mm). Mobile phase A consisted of 0.1% formic acid in water, and mobile phase B contained 0.1% formic acid in acetonitrile. Peptides were separated using a linear gradient from 6% to 25% B over 173 min, ramped to 40% B over 20 min, and then to 90% B over 5 min, at a flow rate of 300 nL/min and column temperature of 35°C. Column regeneration and up to three blank injections were performed between sample runs.

Data were acquired in data-dependent acquisition (DDA) mode. Full MS scans were collected from 350–1,500 m/z at a resolution of 120,000 with an automatic gain control (AGC) target of 3 × 10⁶. MS/MS spectra were acquired for the top 20 precursors per cycle using higher-energy collisional dissociation (HCD) at a normalized collision energy of 30%, with a 1.4 Th isolation window, AGC target of 1 × 10⁵, and maximum injection time of 50 ms. Fragment ions were detected at a resolution of 30,000. Approximately 0.3 µg of peptide was loaded per injection. Data were analyzed using Proteome Discoverer software v2.5 (Thermo Scientific) for label-free quantification (LFQ). The analysis involved two main workflows: a processing workflow for protein identification and a consensus workflow for LFQ-based quantification. Protein identification was performed using the SEQUEST HT search engine against the SwissProt Homo sapiens reference proteome (20,385 sequences). Search parameters included tryptic digestion with up to two missed cleavages, a precursor mass tolerance of 10 ppm, and a fragment mass tolerance of 0.02 Da. Dynamic modifications included methionine oxidation and carbamidomethylation of cysteine. Both target and decoy database searches were conducted, and peptide-spectrum matches were filtered at a 95% confidence level (*P* < 0.05).

LFQ analysis was performed using .msf files generated from the processing workflow. A Feature Mapper node was used for chromatographic alignment of quantifiable features, followed by a Precursor Ion Quantifier node configured to normalize protein abundances based on total peptide amount. Only proteins with a FDR ≤1% and supported by at least two unique peptides were retained for downstream analysis.

DAP analysis was performed using g:ProfileR^50^. Proteins with a FDR < 0.01 and a log₂(fold change) > 1 or < –1 were considered significantly differentially abundant.

### Statistical analysis

For comparative analyses, statistical significance was determined using a two-tailed Student’s t-test or Welch’s t-test, as appropriate. Effect sizes were calculated using Cohen’s *d* to assess the magnitude of group differences. For correlation analyses, Pearson’s correlation test was used (GraphPad Prism v10.3.1). Details of the statistical tests and the number of biological replicates (*n*) for each experiment are provided in the corresponding figure legends.

## Acknowledgements

We acknowledge the invaluable contributions of the patients and families involved in this study. We thank members of Park lab for their support and advice for the project. iPSCs were generated by the Yale hESC/iPSC Core, whose support is gratefully acknowledged. We are grateful to the Center for Cellular and Molecular Imaging and the Electron Microscopy Facility at Yale School of Medicine for their imaging support and technical assistance. We thank the Yale Center for Genome Analysis (YCGA) for Next-generation sequencing. We also thank the MS & Proteomics Resource at Yale University for providing the necessary mass spectrometers and associated biotechnology tools. Computation time was provided by Yale University Biomedical High Performance Computing Center. This study is supported by BD^2^: Breakthrough Discoveries for Thriving with Bipolar Disorder (#DG230102), and National Institutes of Health (R01MH118344). M.S.C. (No. RS-2024-00359456), J.K. (2021R1A6A3A14043871) and W.S.Y. (2021R1A6A3A14043824) are supported by National Research Foundation of Korea. The YCGA is supported by the National Institute of General Medical Sciences of the National Institutes of Health under Award Number 1S10OD030363-01A1. The MS & Proteomics Resource is funded in part by Yale School of Medicine and the Office of the Director, National Institutes of Health (S10OD02365101A1, S10OD019967, and S10OD018034).

## Author information

These authors contributed equally: Jonghun Kim, Mu Seog Choe.

## Contributions

J.K., M.S.C., and I.H.P. conceived the study and wrote the manuscript. J.K. designed the experiments, generated hCOs and hChPOs from all iPSC lines, performed organoid analyses, and analyzed the MS data. M.S.C. designed the experiments and carried out all bioinformatic analyses, including processing and interpretation of transcriptomic data. K.K. analyzed MRI data to quantify neuroanatomical volumes. Y.H. conducted the GWAS analysis. K.N. performed the analysis of WGS data. W.S.Y., F.R.K., C.L., M.S., and H.J.C. provided experimental support. M.L., J.R.G. supervised the genetic analyses. E.A.J. supervised experimental procedures. H.P.B. established the BD cohort, acquired the neuroimaging data, analyzed the MRI data and supervised experimental work. I.H.P led the project. All authors participated in critical revisions to the manuscript. Corresponding authors Correspondence to In-Hyun Park

## Supplementary information

Supplementary Tables 1-10

---

**Extended Data Fig. 1.**
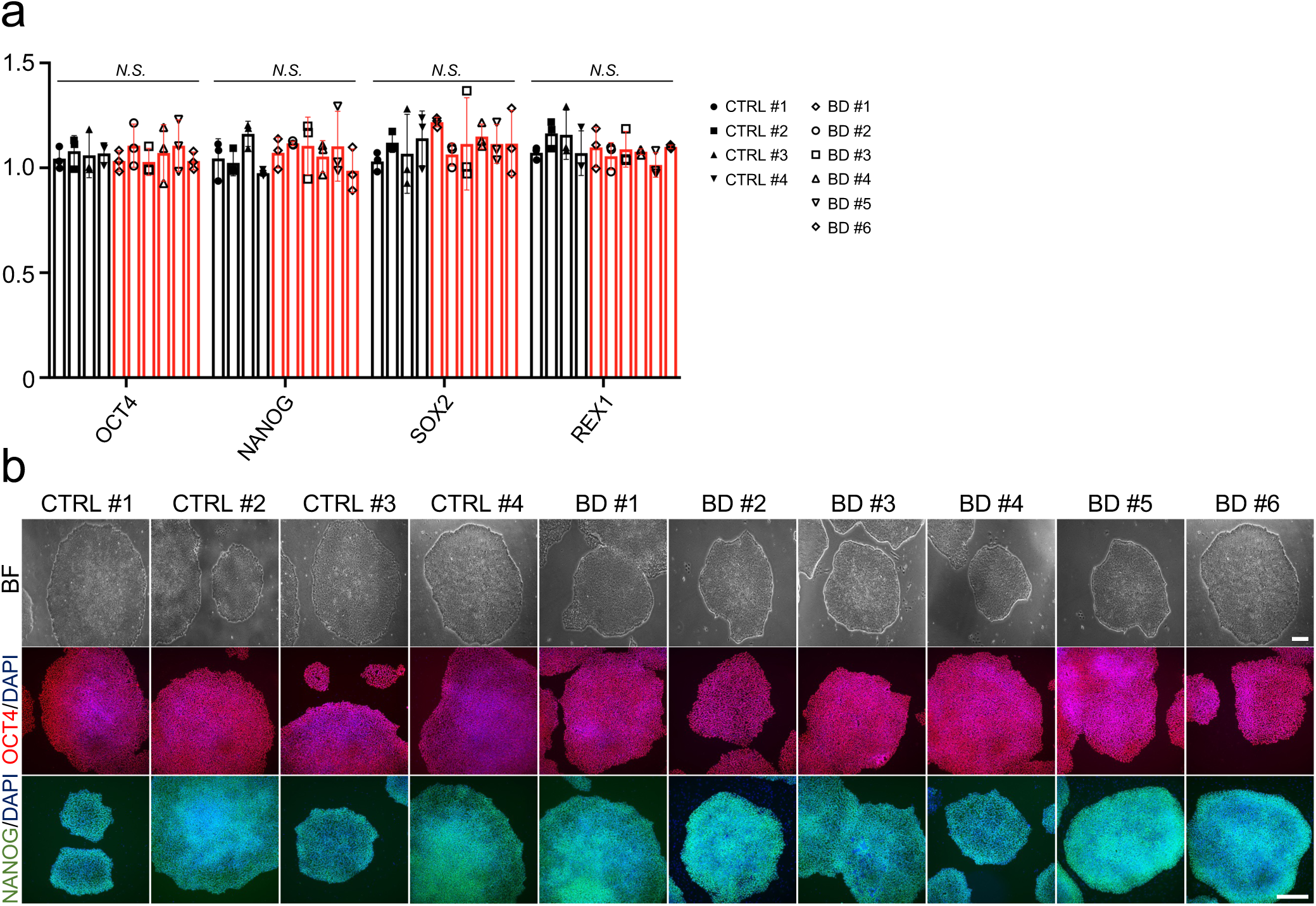
Characterization of control and BD iPSC lines. **a**, Gene expression levels of pluripotency markers across all iPSC lines, confirmed by RT–qPCR (*n* = 3 per sample). **b**, Bright-field morphology (top) and immunofluorescence staining for pluripotency markers (middle and bottom) across all iPSC lines. Scale bars, 100 µm. All data are presented as mean ± s.d. Statistical analysis was performed using a two-tailed Student’s t-test (b). *N.S.*, not significant.

**Extended Data Fig. 2.**
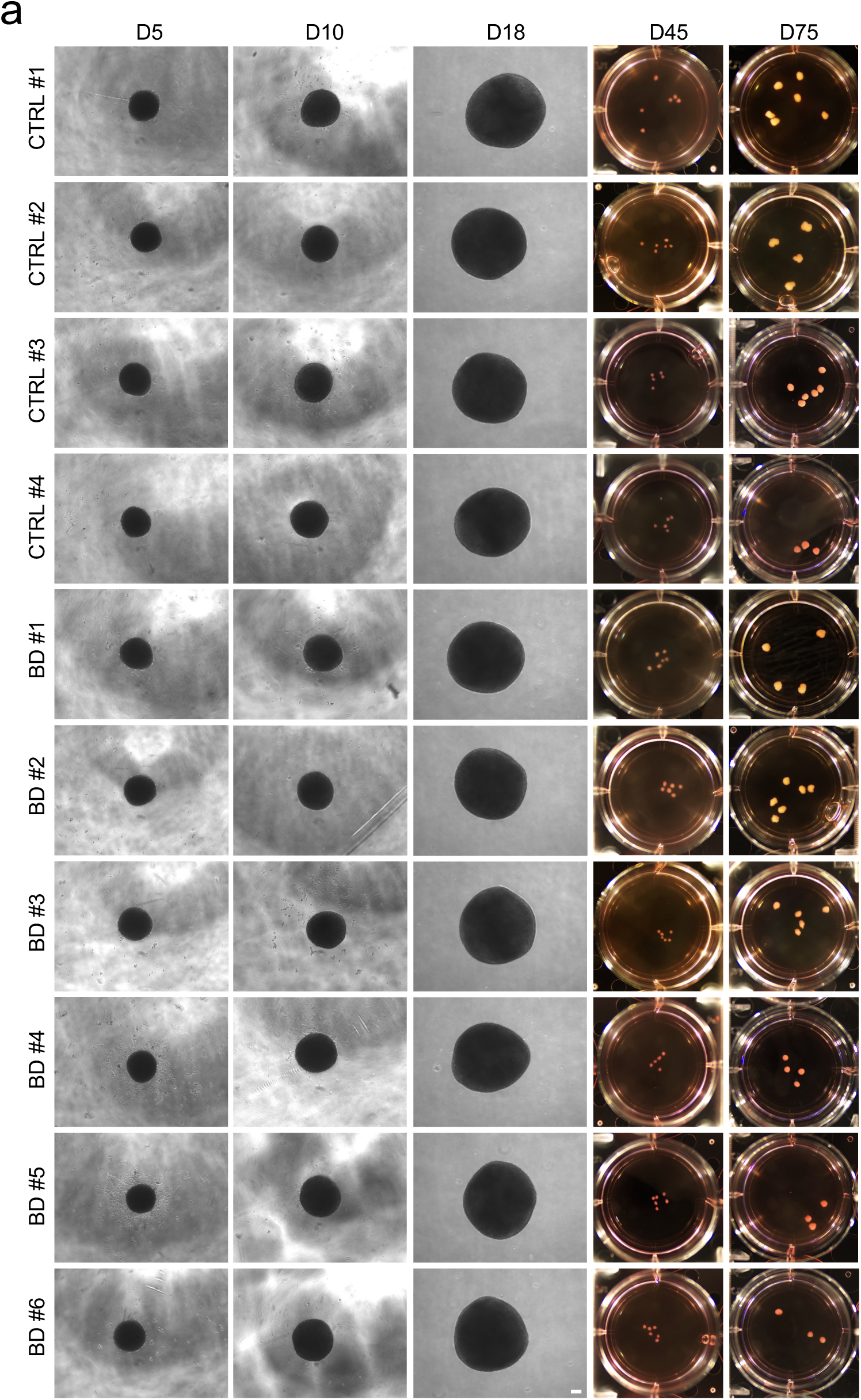
Generation of hCOs from control and BD iPSC lines. **a**, Representative images of hCOs at different time points (Day 5, 10, 18, 45, and 75). Scale bars, 100 µm.

**Extended Data Fig. 3.**
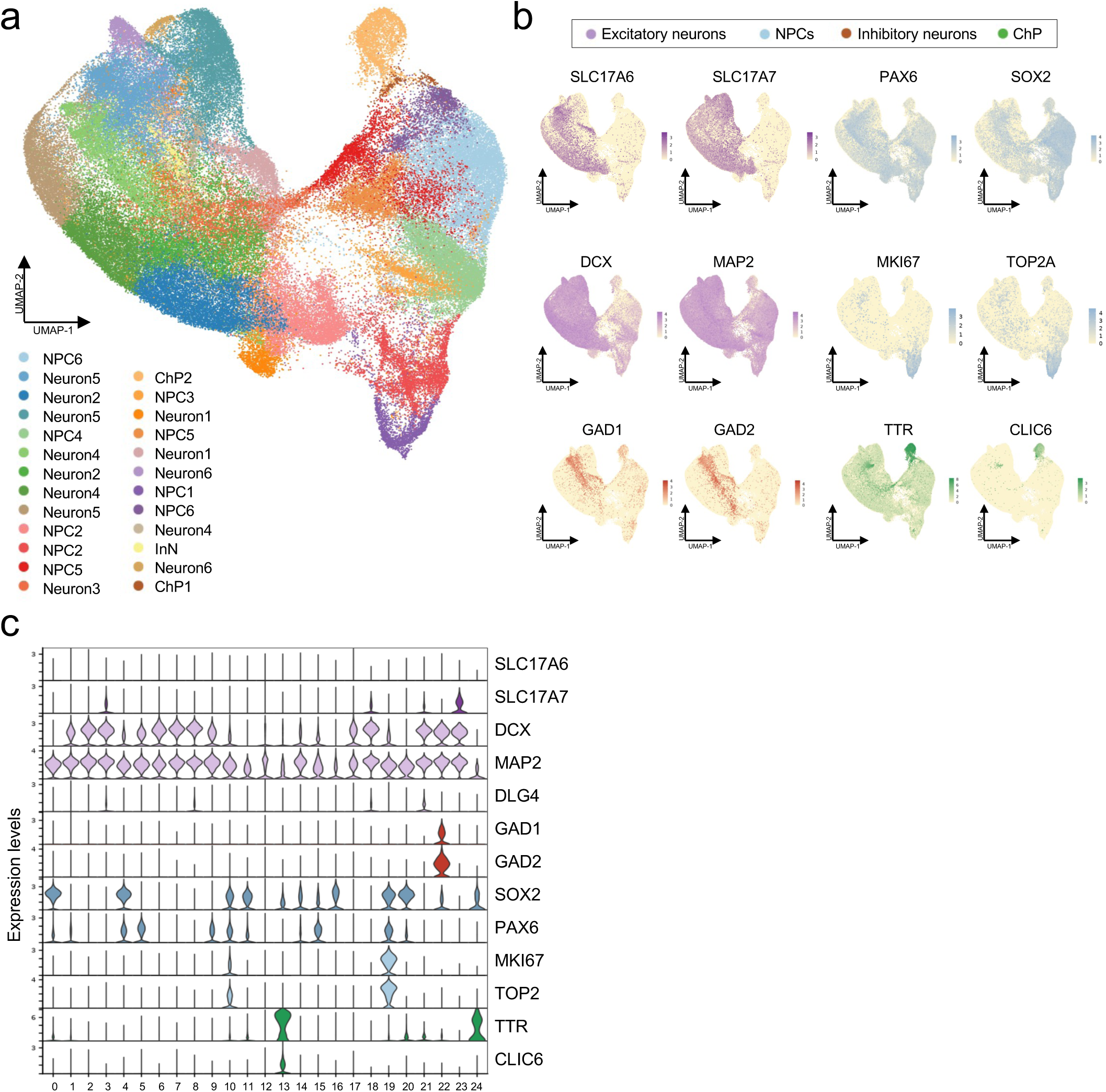
Cluster annotation of cell types identified by scRNA-seq. **a,** UMAP visualization of scRNA-seq data from four control and six BD hCOs, showing 25 distinct cell clusters. **b**, Feature plots showing expression of representative marker genes across distinct cell types. SLC17A6/SLC17A7 for excitatory neurons; DCX/MAP2 for neurons; GAD1/GAD2 for inhibitory neurons; PAX6/SOX2/MKI67/TOP2A for NPCs; TTR/CLIC6 for ChP. **c**, Expression levels of specific marker genes across clusters. NPC, neural progenitor cell; InN, inhibitory neuron; ChP, choroid plexus.

**Extended Data Fig. 4.**
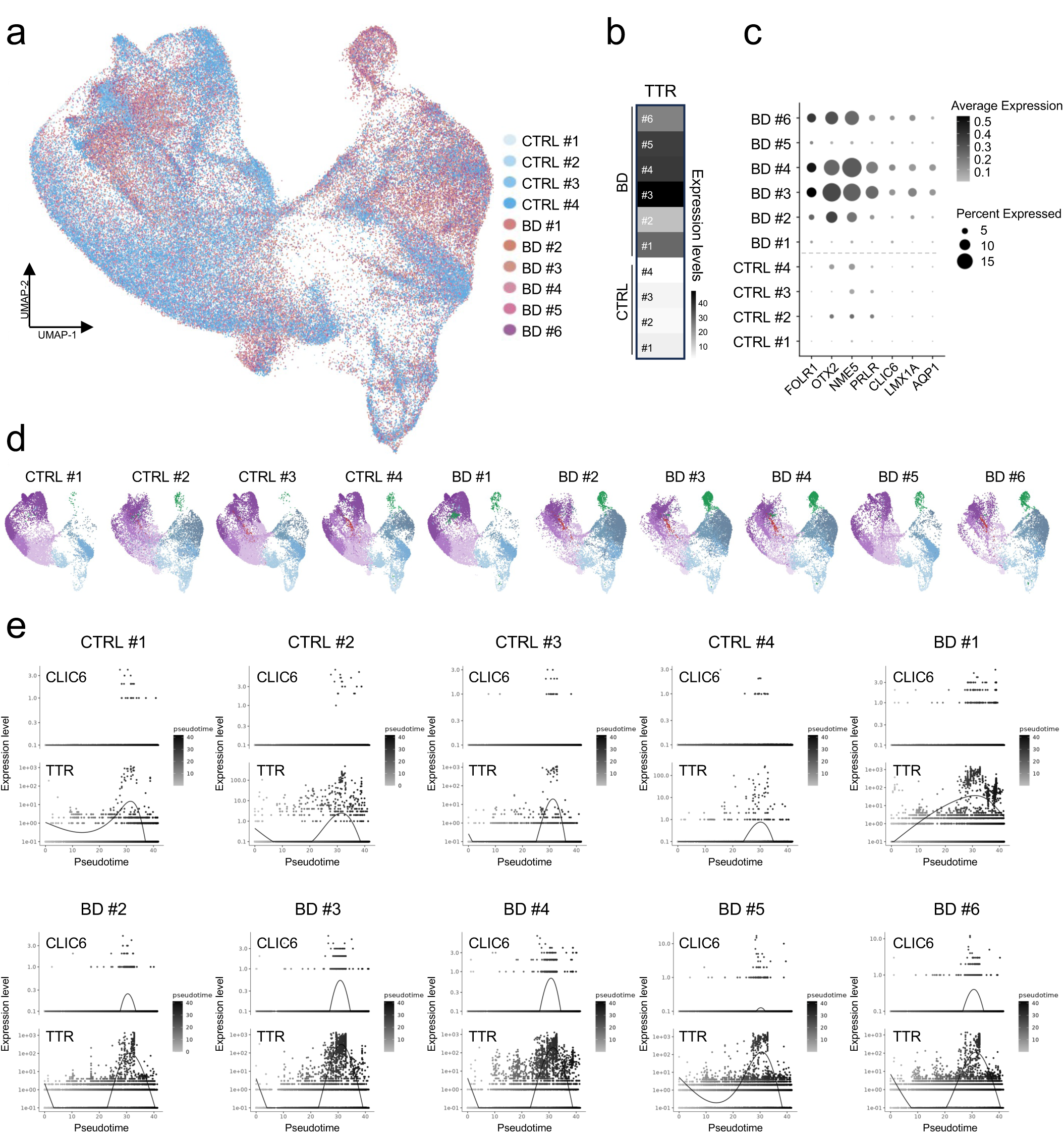
scRNA-seq analysis of individual hCOs. **a**, UMAP showing overlaid cellular distributions from individual hCOs. Blue tones represent control samples, and red tones represent BD samples. **b**, Heatmap exhibiting TTR expression levels across individual hCOs. **c**, Dot plot showing expression of ChP-specific marker genes. **d**, UMAPs of individual hCOs, displayed separately. **e**, Temporal expression dynamics of ChP markers (TTR, CLIC6) along pseudotime across individual hCOs.

**Extended Data Fig. 5.**
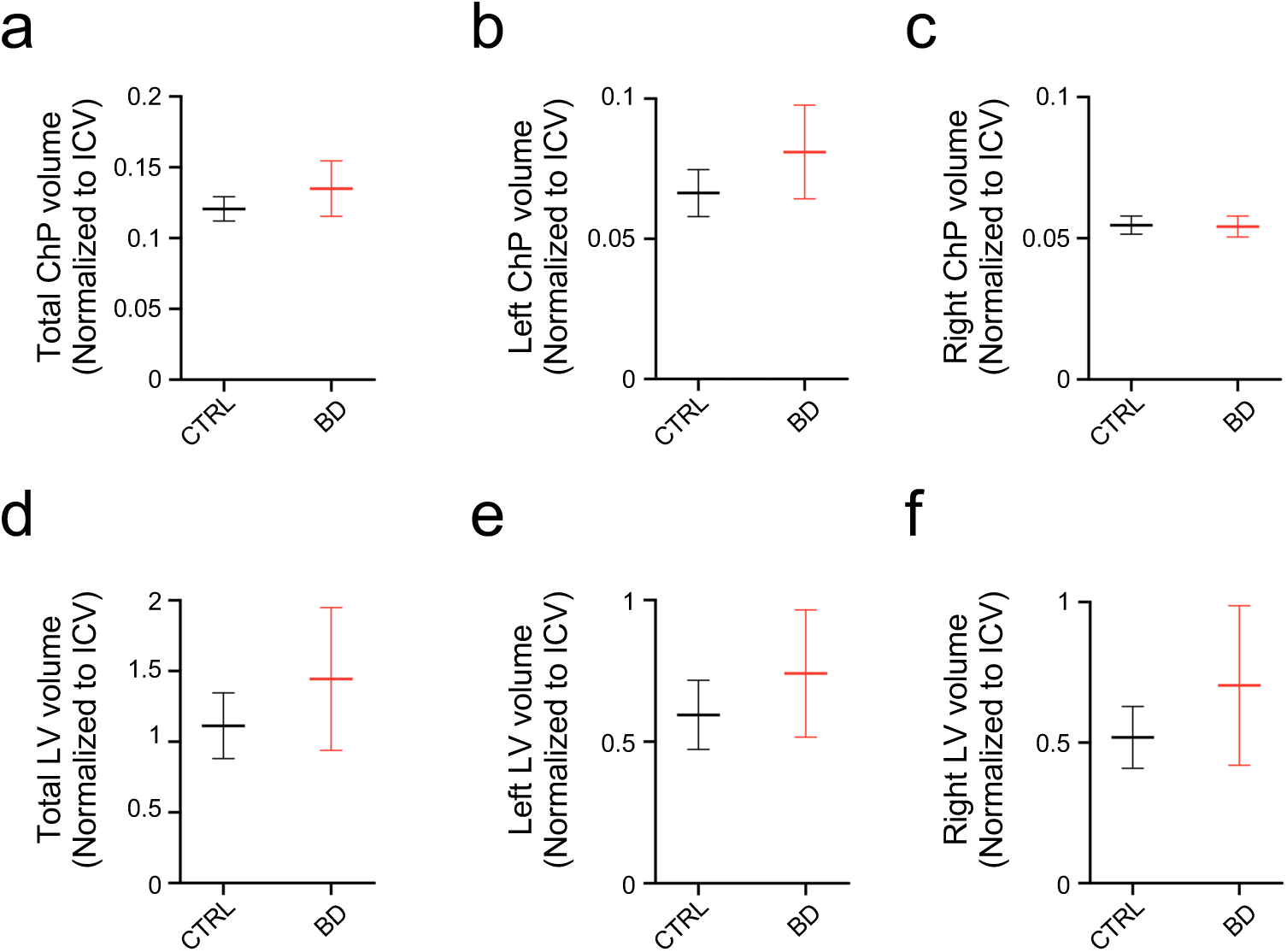
Individuals with BD exhibit ChP enlargement on MRI. **a-f,** Neuroanatomical volume measurements in controls (*n* = 3) and individuals with BD (*n* = 6). (a) Total ChP volume. (b) Left ChP volume. (c) Right ChP volume. (d) Total LV volume. (e) Left LV volume. (f) Right LV volume. All data are presented as mean ± s.e.m. Statistical analysis was performed using a two-tailed Welch’s t-test (a-f) and Cohen’s *d* for effect size (a-f). All comparisons were not significant. Effect sizes: total ChP volume = 0.4, left ChP volume = 0.49, right ChP volume = 0.14, total LV volume = 0.36, left LV volume = 0.35, right LV volume = 0.36. LV, lateral ventricle; ICV, intracranial volume.

**Extended Data Fig. 6.**
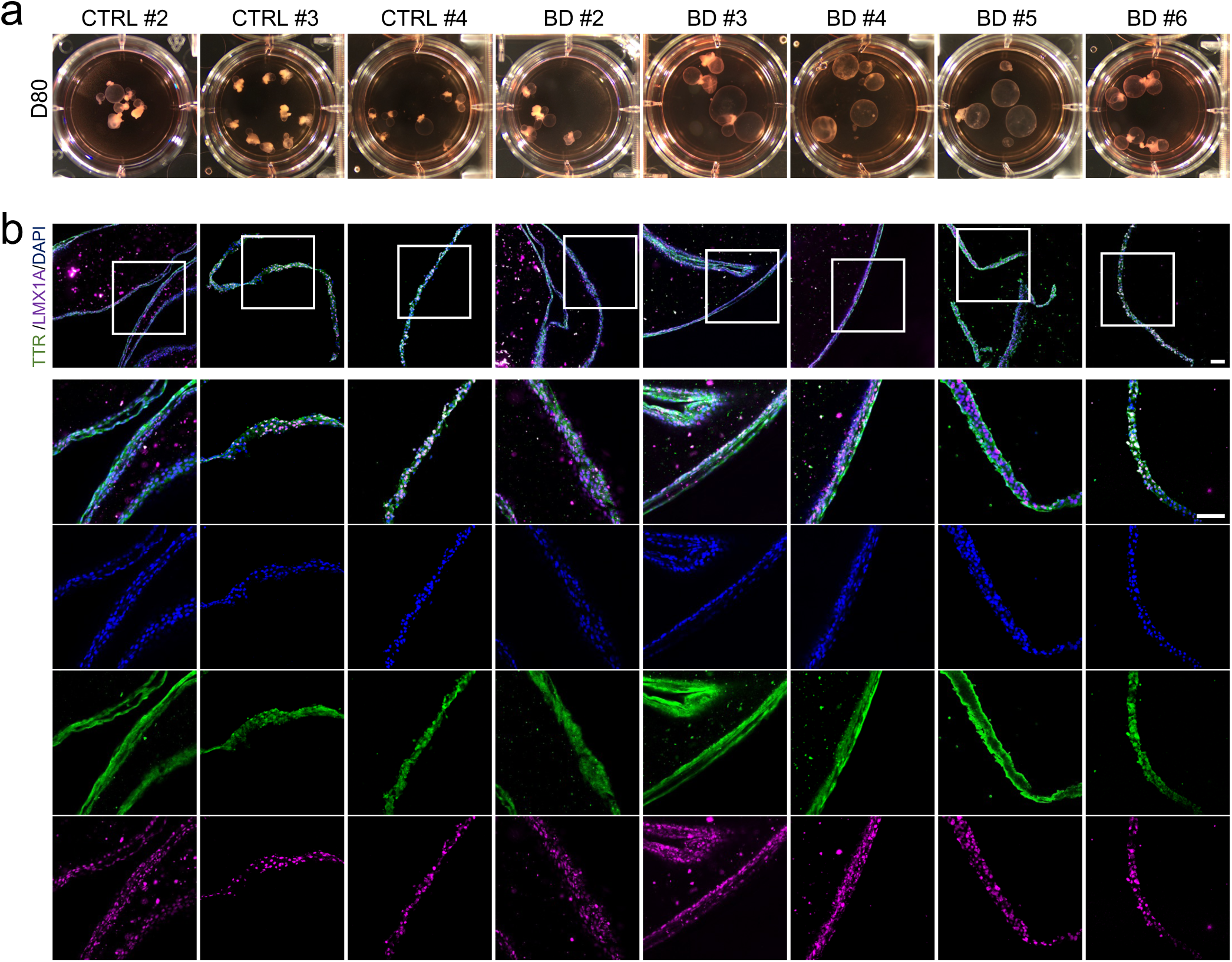
Generation and characterization of hChPOs. **a,** Representative images of hChPOs at Day 80 from both control and BD groups. **b**, Immunofluorescence images of TTR and LMX1A expression in all hChPOs at Day 80. Nuclei are counterstained with DAPI. The white box indicates the magnified region shown below. Scale bars, 100 µm.

**Extended Data Fig. 7.**
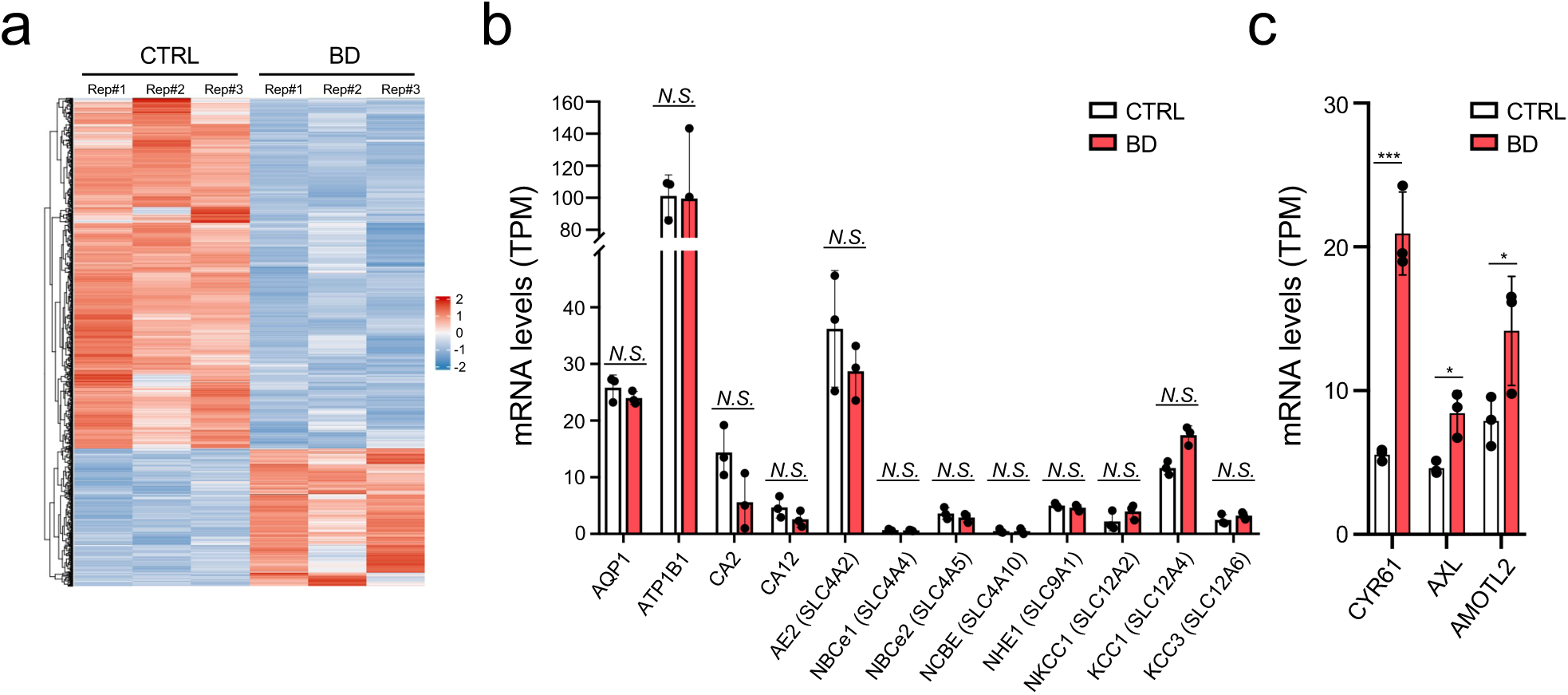
Transcriptomic analysis of control and BD hChPOs. **a**, Heatmap showing distinct gene expression patterns between control and BD hChPOs (*n* = 3 per group). **b**, Expression levels of ion channel and transporter genes involved in CSF secretion, based on TPM values (*n* = 3 per group). **c**, Expression levels of YAP/TAZ target genes, based on TPM values (*n* = 3 per group). All data are presented as mean ± s.d. Statistical analysis was performed using a two-tailed Student’s t-test (b,c). \*\*\**P* < 0.001, \**P* < 0.05, *N.S*., not significant.

**Extended Data Fig. 8.**
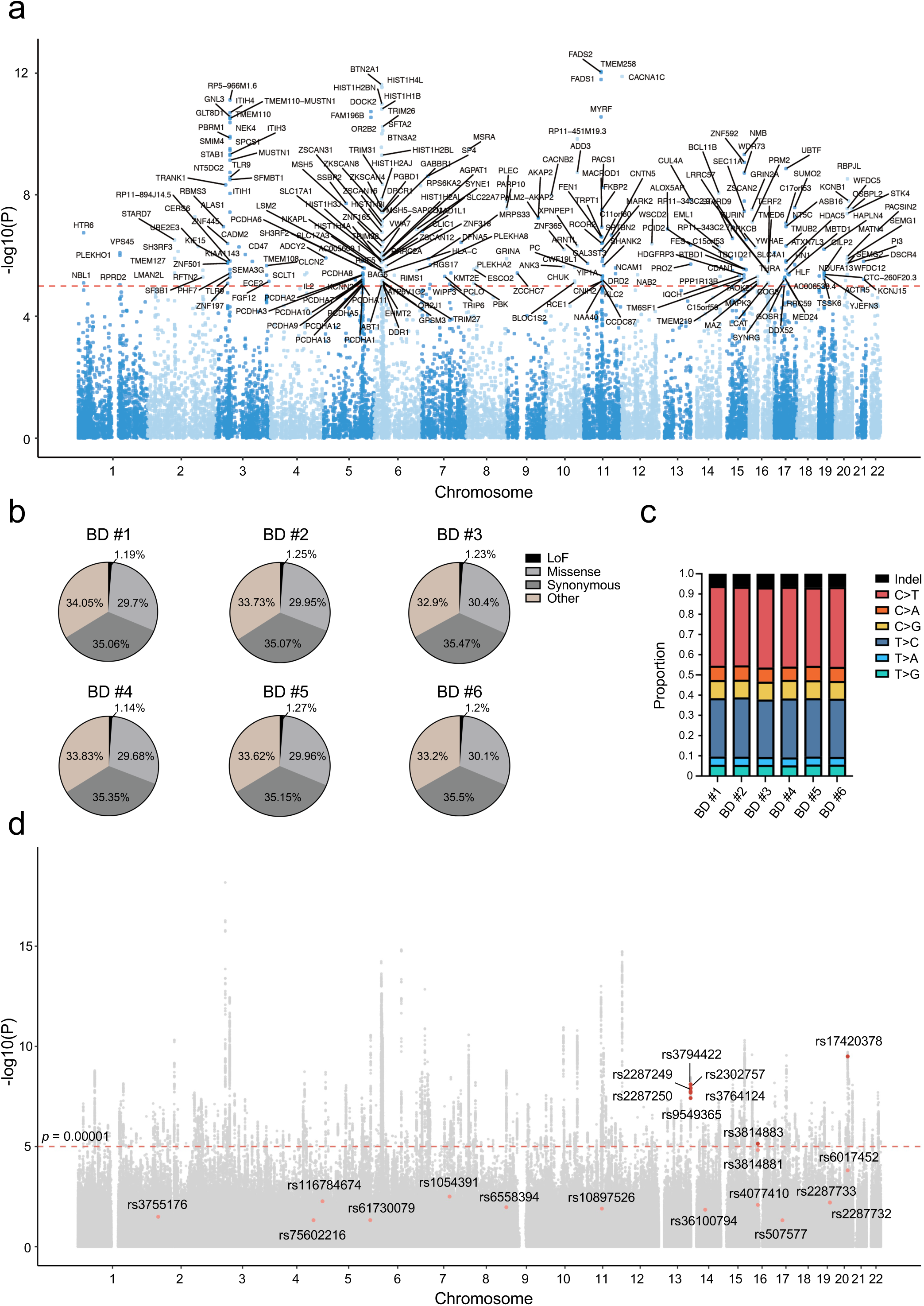
Genetic analysis of controls and individuals with BD. **a**, Gene-based Manhattan plot summarizing genetic associations from GWAS in a large BD cohort (*n* = 371,549 controls; *n* = 41,917 BD), with gene labels shown only for associations at *P* < 1 × 10⁻⁵. **b**, Pie charts showing the proportion of mutation types across six individuals with BD, including loss-of-function (LoF), missense, synonymous, and other variants. **c**, Bar charts illustrating the distribution of indel and SNV substitution types across six individuals with BD. **d**, SNP-based Manhattan plot highlighting significant SNPs in genes associated with the Hippo signaling pathway. Red and pink dots indicate associations at *P* < 1 × 10⁻⁵ and *P* < 0.05, respectively. Indel, insertion or deletion; SNV, single nucleotide variant.

**Extended Data Fig. 9.**
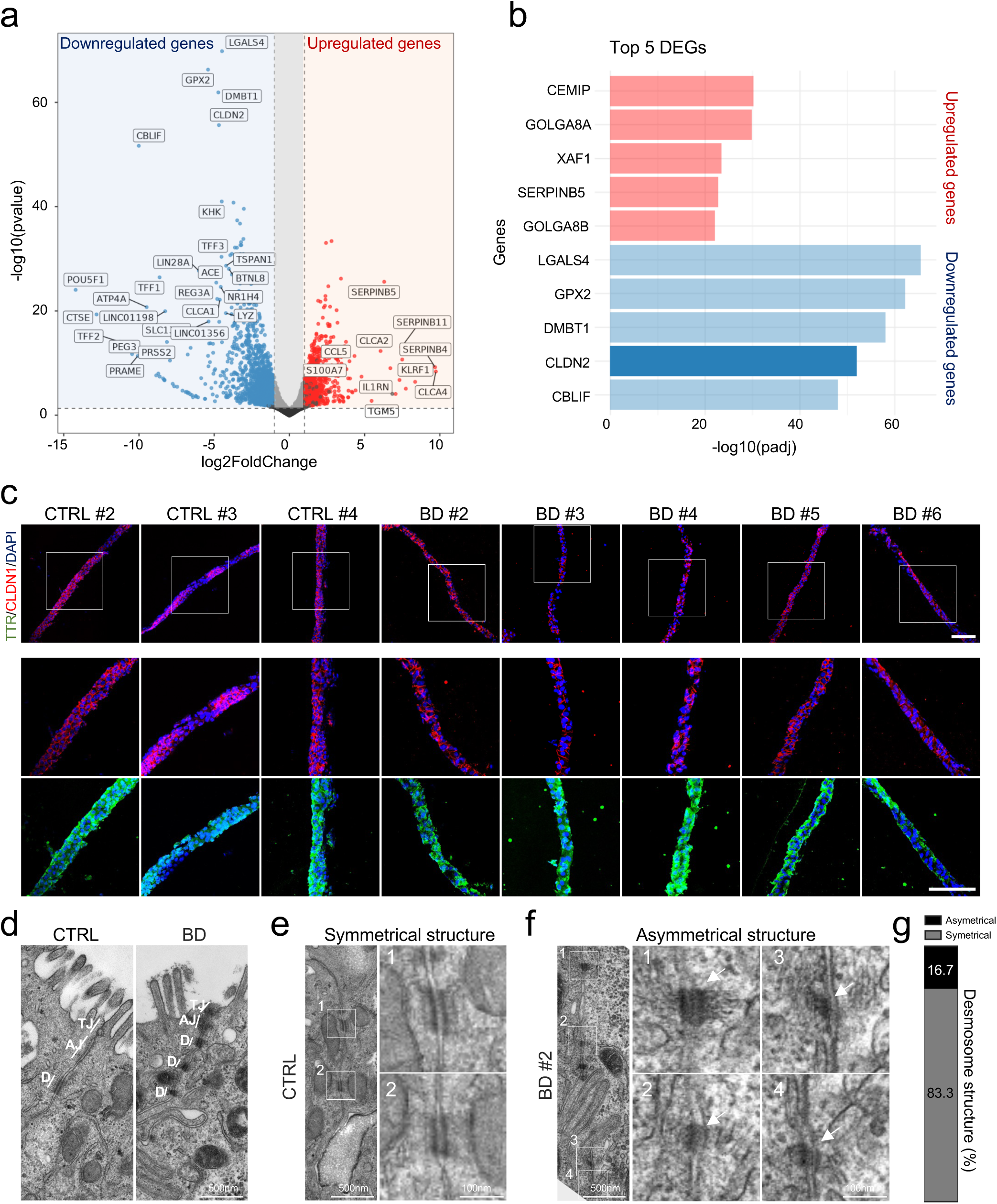
Analysis of cell junctions in control and BD hChPOs. **a**, DEGs between control and BD hChPOs. Red dots indicate upregulated genes (log_2_ fold change > 1, FDR < 0.05), and blue dots indicate downregulated genes (log_2_ fold change < -1, FDR < 0.05) in BD hChPOs. **b**, Top five DEGs between control and BD hChPOs. Red bars represent upregulated genes in BD hChPOs, while blue bars represent downregulated genes. The dark blue bar highlights *CLDN2*, a gene associated with cell junctions. **c**, Immunofluorescence images of TTR and CLDN1 expression in all hChPOs at Day 80. Nuclei are counterstained with DAPI. The white box indicates the magnified region shown below. Scale bars, 100 µm. **d**, Ultrastructural images of cell junctions in control and BD hChPOs. Scale bars, 500 nm. **e**, Ultrastructural images showing the symmetrical structure of desmosomes in control hChPOs. White boxes indicate the magnified regions shown in panels 1 and 2. Scale bars, 500 nm (left), 100 nm (right). **f**, Ultrastructural images showing the asymmetrical structure of desmosomes in BD hChPOs from a different individual (BD #2). White boxes indicate the magnified regions shown in panels 1–4. The white arrow highlights the asymmetric structure of the desmosome in the high-magnification image. Scale bars, 500 nm (main), 100 nm (1–4). **g**, Quantification of desmosome structural distributions shown in f (*n* = 7). TJ, tight junction; AJ, adherens junction; D, desmosome.

**Extended Data Fig. 10.**
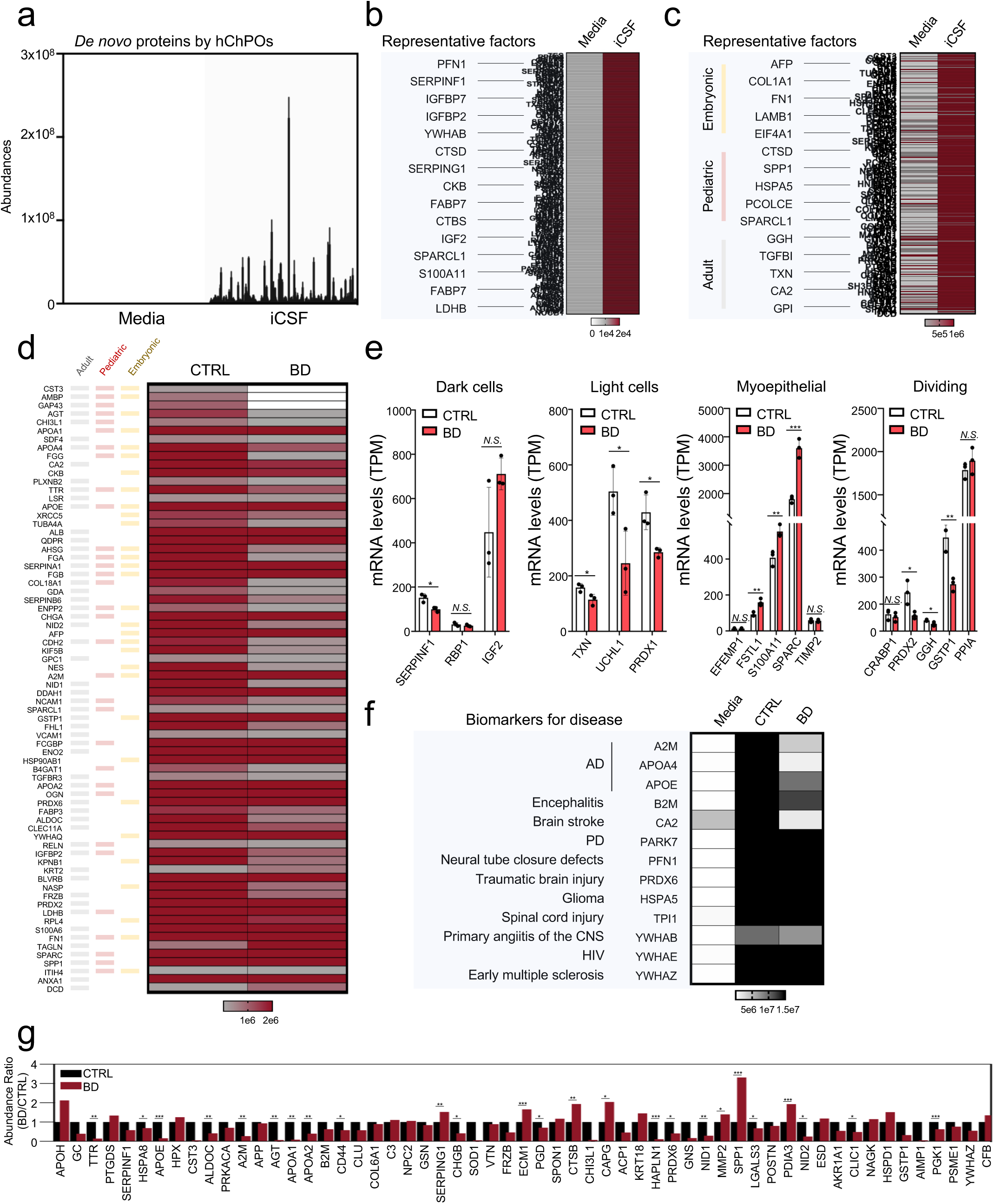
Proteomic analysis of control and BD hChPOs. **a**, Detection of de novo proteins in hChPO-derived iCSF, but not present in the media. **b**, Heatmap displaying the secretion profiles of de novo proteins identified in hChPO-derived iCSF. Representative proteins are highlighted in the blue box. **c**, Heatmap showing the secretion profiles of developmental stage-specific proteins identified in hChPO-derived iCSF. Representative proteins are highlighted in the blue box. **d**, Heatmap exhibiting the secretion profiles of developmental stage-specific proteins among DAPs between control and BD iCSF. **e**, Expression levels of ChP epithelial cell type-specific genes, based on TPM values (*n* = 3 per group). **f**, Heatmap showing disease-associated biomarkers identified in hChPO-derived iCSF. Proteins and their related diseases are highlighted in the blue box. **g**, Comparative abundance BD-associated proteins in control and BD iCSF (*n* = 3 per group). All data are presented as mean ± s.d. Statistical analysis was performed using a two-tailed Student’s t-test (e,g). \*\*\**P* < 0.001, \*\**P* < 0.01, \**P* < 0.05, *N.S*., not significant.

